# Dynamic transcriptome profiles within spermatogonial and spermatocyte populations during postnatal testis maturation revealed by single-cell sequencing

**DOI:** 10.1101/464149

**Authors:** Kathryn J. Grive, Yang Hu, Eileen Shu, Andrew Grimson, Olivier Elemento, Jennifer K. Grenier, Paula E. Cohen

**Author notes:** Corresponding Authors: J.K. Grenier and P.E. Cohen.

## Abstract

Spermatogenesis is the process by which male gametes are formed from a self-renewing population of spermatogonial stem cells (SSCs) residing in the testis. SSCs represent less than 1% of the total testicular cell population, but must achieve a stable balance between self-renewal and differentiation. Once differentiation has occurred, the newly formed and highly proliferative spermatogonia must then enter the meiotic program in which DNA content is doubled, then halved twice to create haploid gametes. While much is known about the critical cellular processes that take place during the specialized cell division that is meiosis, much less is known about how the spermatocytes in the “first-wave” compare to those that contribute to long-term, “steady-state” spermatogenesis. Given the strictly-defined developmental process of spermatogenesis, this study was aimed at exploring the transcriptional profiles of developmental cell stages over the age of the animal. Using a combination of comprehensive germ cell sampling with high-resolution, single-cell-mRNA-sequencing, we have generated a reference dataset of germ cell gene expression. We show that discrete developmental stages possess significant differences in the transcriptional profiles from neonates compared to juveniles and adults. Importantly, these gene expression dynamics are also reflected at the protein level in their respective cell types. We also show differential utilization of many biological pathways with age in both spermatogonia and spermatocytes, demonstrating significantly different underlying gene regulatory programs in these cell types over the course of testis development and spermatogenic waves. This dataset represents the first unbiased sampling of spermatogonia and spermatocytes in the developing testis over developmental age, at high-resolution, single-cell depth. Not only does this analysis reveal previously unknown transcriptional dynamics of a highly transitional cell population, it has also begun to reveal critical differences in biological pathway utilization in developing spermatogonia and spermatocytes, including response to DNA damage and double-strand breaks.

**Author Summary:** Spermatogenesis is the process by which male gametes – mature spermatozoa – are produced in the testis. This process requires exquisite control over many developmental transitions, including the self-renewal of the germline stem cell population, commitment to meiosis, and ultimately, spermiogenesis. While much is known about molecular mechanisms regulating single transitions at single time points in the mouse, much less is understood about how the spermatogenic progenitor cells, spermatogonia, or the meiotic cells, spermatocytes, of the testis change over developmental age.

Our single-cell-mRNA-sequencing analysis is the first to profile both spermatogonia and spermatocytes from neonatal mice through adulthood, revealing novel gene expression dynamics and differential utilization of biological pathways. These discoveries help us to understand how the spermatogenic progenitors of this population modulate their activity to adapt to a changing testicular environment. Furthermore, they also begin to explain previously-observed differences - and deficiencies - in spermatocytes that are derived from the first “wave” of spermatogenesis. Overall, this dataset is the first of its kind to comprehensively profile gene expression dynamics in male germ cell populations over time, enriching our understanding of the complex and highly-orchestrated process of spermatogenesis.

## Introduction

Mammalian spermatogenesis requires proper establishment of the spermatogonial stem cell (SSC) pool, which resides within the seminiferous tubules of testis and supports life-long germ cell development^1^. These progenitors give rise to all the differentiating germ cells of the mouse testis, ranging from spermatogonia to spermatocytes to spermatids, and finally to mature spermatozoa. Despite the essential nature of this process, the genetic regulatory mechanisms underlying the many complex cellular transitions, and the maturation of this system during testis development, have yet to be fully described.

Gamete development in the mouse relies on a rare population of primordial germ cells, the bi-potential progenitors of all gametes, which are specified in the developing embryo at embryonic day (E) 6.25 ^2^. These cells migrate to and colonize the developing gonad, arriving at the genital ridge from E10.5 ^3^, and undergo abundant proliferation until E13.5. At this time, germ cells developing in an XX (female) gonad will enter the meiotic program as oocytes, while germ cells developing in an XY (male) gonad will become prospermatogonia, remaining relatively non-proliferative until shortly after birth^4,5^. Prospermatogonia are able to adopt several fates^6^: in early postnatal life, a subset of these cells differentiate immediately into spermatogonia and continue to progress through spermatogenesis, to constitute the “first wave” of spermatogenesis. A second subset of prospermatogonia will undergo apoptosis, while the remaining prospermatogonia will become established within the testicular stem cell niche soon after birth, to become the self-renewing SSC population that will support “steady-state” spermatogenesis throughout life. This germline stem cell population makes up less than 1% of the cells of adult testes^7^, and must balance self-renewal and differentiation to maintain a healthy male gamete supply. Thus, the first cohort of meiotically-active male germ cells enter the meiotic program without first entering a self-renewal SSC phase, clearly differentiating the first wave of spermatogenesis from the other subsequent waves.

SSCs are triggered to enter spermatogenesis coincident with a burst of retinoic acid (RA), which induces both the spermatogonial divisions and the entry into prophase I of meiosis^8–11^. Thus, in mice, male meiotic entry commences around postnatal day (PND) 10, in response to RA-induced expression of key genes, including ‘Stimulated by Retinoic Acid 8’ (*Stra8*)^8–11^. Spermatocytes execute many essential meiotic events including creation of double-strand breaks, synapsis of homologous chromosomes, and DNA repair and crossover formation, all of which are critical to proper segregation of homologs in the first meiotic division. Failure to properly execute any of these steps is known to result in potential chromosome mis-segregation, non-disjunction events, aneuploidy, and infertility (reviewed comprehensively in ^12,13^).

While the developmental transitions which underlie germ cell differentiation and maturation have been broadly defined, the gene regulatory underpinnings of these transitions remain largely uncharacterized. Furthermore, studies which have shed some light upon genetic regulatory mechanisms of these processes often focus on single time points, or utilize cell enrichment protocols that may bias the output. In this manuscript, we have performed the first single-cell sequencing developmental time series of the male mouse germline with comprehensive/unbiased sampling, thereby capturing all germ cell types through the progression of postnatal testis maturation. The advent of single cell transcriptomics provides an invaluable tool for understanding gene expression dynamics at very high resolution in a large number of individual cells in parallel. Furthermore, single-cell sequencing reveals heterogeneity and potential plasticity within cell populations, which bulk mRNA sequencing is unable to accomplish, making it an ideal tool for profiling germ cell populations which rapidly progress through myriad developmental transitions.

We demonstrate that germ cells display novel gene regulatory signatures over time, while cells positive for single protein markers have the capacity to change dramatically with age, and therefore cells of a particular “identity” may differ significantly from postnatal to adult life. Intriguingly, we have also begun to identify differential expression of genes in critical biological pathways which may contribute to observed differences in the first-wave of spermatogenesis^14,15^. Dissecting the complex dynamics of these developmental transitions can provide critical information about the transcriptional landscape of both SSCs, spermatogonia, and spermatocytes, and the regulatory mechanisms that underlie the formation of a dynamic and functional complement of germ cells to support life-long spermatogenesis.

## Results

### Single-cell sequencing from testes of different developmental ages robustly defines germ cell populations

Mouse testes were collected at several time points, selected to represent distinct stages of germline development: postnatal day (PND) 6 (during SSC specification), PND 14 (first appearance of pachytene spermatocytes during the first wave), PND 18 (pachytene and diplotene spermatocytes from the first wave present), PND 25 (spermatids present) and PND30 (spermatozoa present) (**Figure 1A**) and subjected to single-cell RNAseq. The tissue was dissociated, and the resulting slurry subjected to 30% Percoll sedimentation to remove debris. The PND 18 cell suspension was split and processed either with or without Percoll sedimentation as a technical control; due to similarities between libraries, the data from these libraries was thereafter combined (**Figure S1**). Additionally, due to the proportionally high representation of sperm in the adult testis, it was necessary to increase representation of other germ cell types from these samples. To accomplish this goal, an adult testis suspension post-Percoll sedimentation was split in half and either positively magnetically-cell-sorted (MACS) for the cell surface marker THY1, in an attempt to enrich for spermatogonia^16^, or negatively MACS-sorted for ACRV1, in an attempt to deplete testicular sperm^17^. While neither strategy can accomplish complete enrichment of spermatogonia or removal of spermatozoa, respectively, both adult libraries had a representative sample of all germ cell types (**Figure 1B**), and are therefore treated as adult replicates in these data. For each single-cell testis suspension, 4-5,000 cells per mouse were processed through the 10X Genomics Chromium System using standard protocols for single cell RNA sequencing. Libraries were sequenced to average depth 98M reads; on average, 91% of reads mapped to the reference genome. After standard data processing, we obtained gene expression profiles for approximately 1,200-2,500 cells per library (**Figure 1B, 2A**) with representation of between 2,500-5,000 genes per cell (**Figure S2**), all of which indicates the robustness of the sequencing method, and the comparability to other single-cell studies.

**Figure 1.**
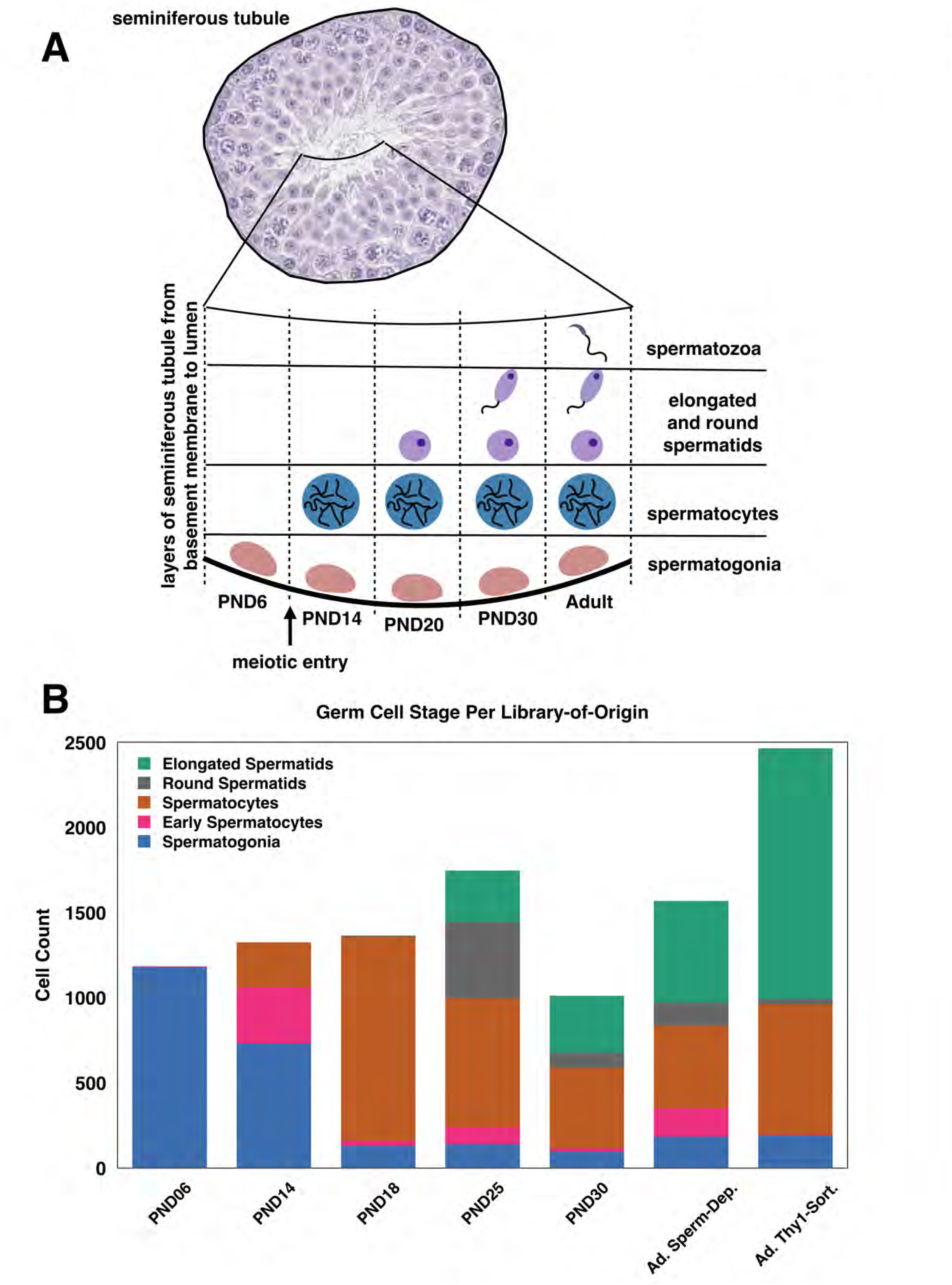
Germ cells profiled in single-cell sequencing analysis are representative of known biology of the developing testis. A) Schematic of the developing testis with germ cell representation at each time point. Only spermatogonia are present during the first week of life, until meiotic entry at PND10, after which germ cells can commit meiosis and progress through spermatogenesis and spermiogenesis, producing the first mature spermatozoa from the first wave around PND30. B) Germ cell composition by proportion and absolute cell number from each library-of-origin.

Primary cell clusters revealed by Seurat (**Figure S3**) were identified and merged into superclusters (**Figure 2B**) based on known marker gene expression (**Table S1**). While somatic cells were evident in the clustering analysis, including Sertoli cells, smooth muscle, and epithelial and hematopoetic cells, they represent a minority (fewer than 25%) of the total cells profiled. In particular, Sertoli cells were primarily derived from the PND6 library (**Figure** 2) due to their increased representation at that time point, as well as processing steps which effectively removed these cells, which become considerably larger in older mice. Therefore, somatic cells were excluded from all additional analysis, which focused on the spermatogonial and spermatocyte populations in the developing testis. Analysis of differentially expressed genes at each time point identifies known marker genes in each cell type, including *Zbtb16* (*Plzf*), *Sall4*, *Sohlh1*, and *Dmrt1* in spermatogonia; *Meioc*, *Prdm3*, *Top2a*, and *Smc3* in early spermatocytes; *Sycp1/2/3* and *H2afx* in spermatocytes; *Acrv1*, *Izumo1*, and *Catsper3/4* in round spermatids; and *Prm3*, *Izumo3*, and *Tssk6* in elongated spermatids (**Figure 3, S4 & S5, Table S1**).

**Figure 2.**
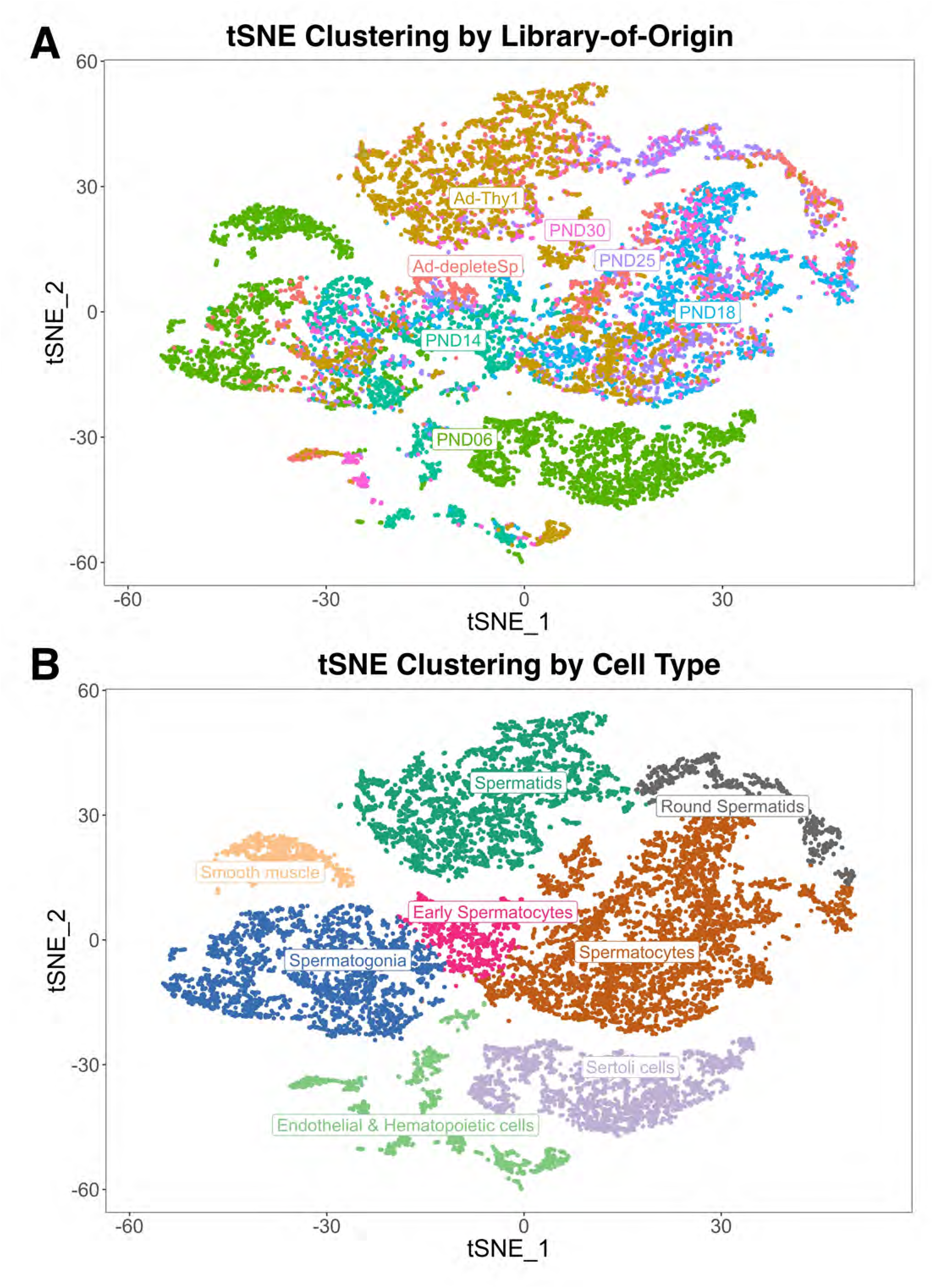
Clustering of single-cell data into libraries-of-origin and cell type classifications. A) tSNE representation of all cells with >500 detected genes and >2000 UMIs, color-coded by library-of-origin B) tSNE representation of all cells with >500 detected genes and >2000 UMIs, color-coded by cell type classification.

**Figure 3.**
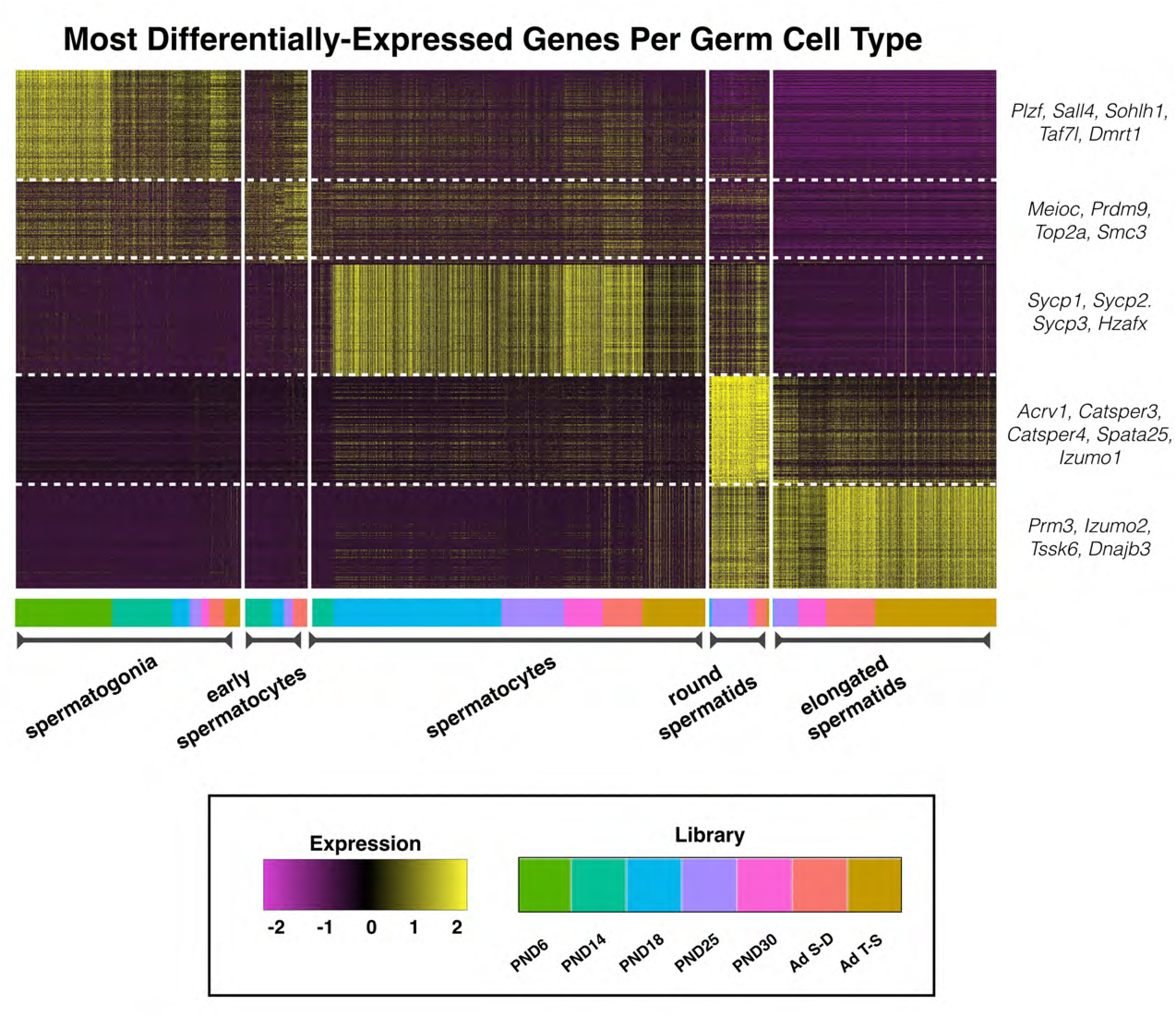
Marker gene heatmap of all germ cell types reveals known signatures. Heatmap of most-differentially-expressed marker genes per germ cell type. Color bar at the bottom indicates library-of-origin time point for cells within each block. Expression is represented as a z-score ranging from −2 to 2. Notable marker genes for each germ cell type are highlighted to the right of the heatmap. The adult sperm-depleted sample is named “Ad S-D” while the adult THY1-sorted sample is named “Ad T-S”.

Critically, the germ cell type classifications are representative of the known timeline of the developing testis (**Figure 1B**), with only spermatogonia present at PND6, some early spermatocytes present at PND14, much greater representation of those spermatocytes at PND18, and appearance of more differentiated round and elongated spermatids from PND25 onwards. Interestingly, we observed the greatest enrichment of spermatids in the positively THY1-sorted adult sample, likely due to non-specific binding of the antibody to the developing acrosome. Despite this, the library contained strong representation of spermatogonia and spermatocytes and was therefore retained in the analysis. The negative ACRV1 sorting for the other adult sample retained representation of all germ cell types in the adult testis, including spermatogonia, which would otherwise have been poorly represented due to the much greater abundance of more differentiated cell types. Overall, both adult samples provide excellent representation for all germ cell types present in the adult testis and are therefore included in this sampling analysis.

Cell-free RNA contamination from lysed cells is a well-known confounding feature in single-cell sequencing libraries, as highly-expressed transcripts from even a small number of lysed cells can become incorporated in the gel bead emulsions of single-cell microfluidics devices^18^. As a result of the incorporation of these transcripts into libraries of cells from which they did not originate, cells which do not endogenously express such transcripts can appear to have low levels of expression of these markers. In this data set, genes highly expressed in elongated spermatids/sperm were detected at low levels in cells identified as spermatogonia and spermatocytes exclusively in libraries made from testes aged 25 days or older (data not shown), the only samples in which spermatids are present. Therefore, we believe the detection of these transcripts in both spermatogonia and spermatocytes of older mice is due to contamination from cell-free RNA derived from lysed spermatids. To mitigate the age-related biases this signal might pose in down-stream analysis, markers of the spermatid/sperm population (genes with a greater than 20:1 ratio of expression between spermatids and other germ cell types), such as *Prm1/2*, have been filtered from the data set (**Table S2**).

### Spermatogonia display characteristic transcriptional signatures, but also novel gene expression dynamics, over developmental age

To better understand the developmental transitions that spermatogonia undergo with age, genes variably-expressed with age were identified by Model-based Analysis of Single Cell Transcriptomics (MAST)^19^ (**Table S3**) and visualized as a heatmap (**Figure 4**). As spermatogonia become proportionally rarer with age (and therefore later-aged individual libraries experience low representation), spermatogonia from libraries PND18, PND25, and PND30 were merged with each other, as were spermatogonia from the two adult samples. While marker genes, indicated in the top row, remain quite constant over time, clear and novel differences can be observed in spermatogonial gene expression over time, particularly in spermatogonia derived from PND6 testes. Several genes, including those noted to the side of the heatmap, are observed to have robust differential expression over time, and may play a role in the establishment and growth of this spermatogonial pool which will support lifelong spermatogenesis.

**Figure 4.**
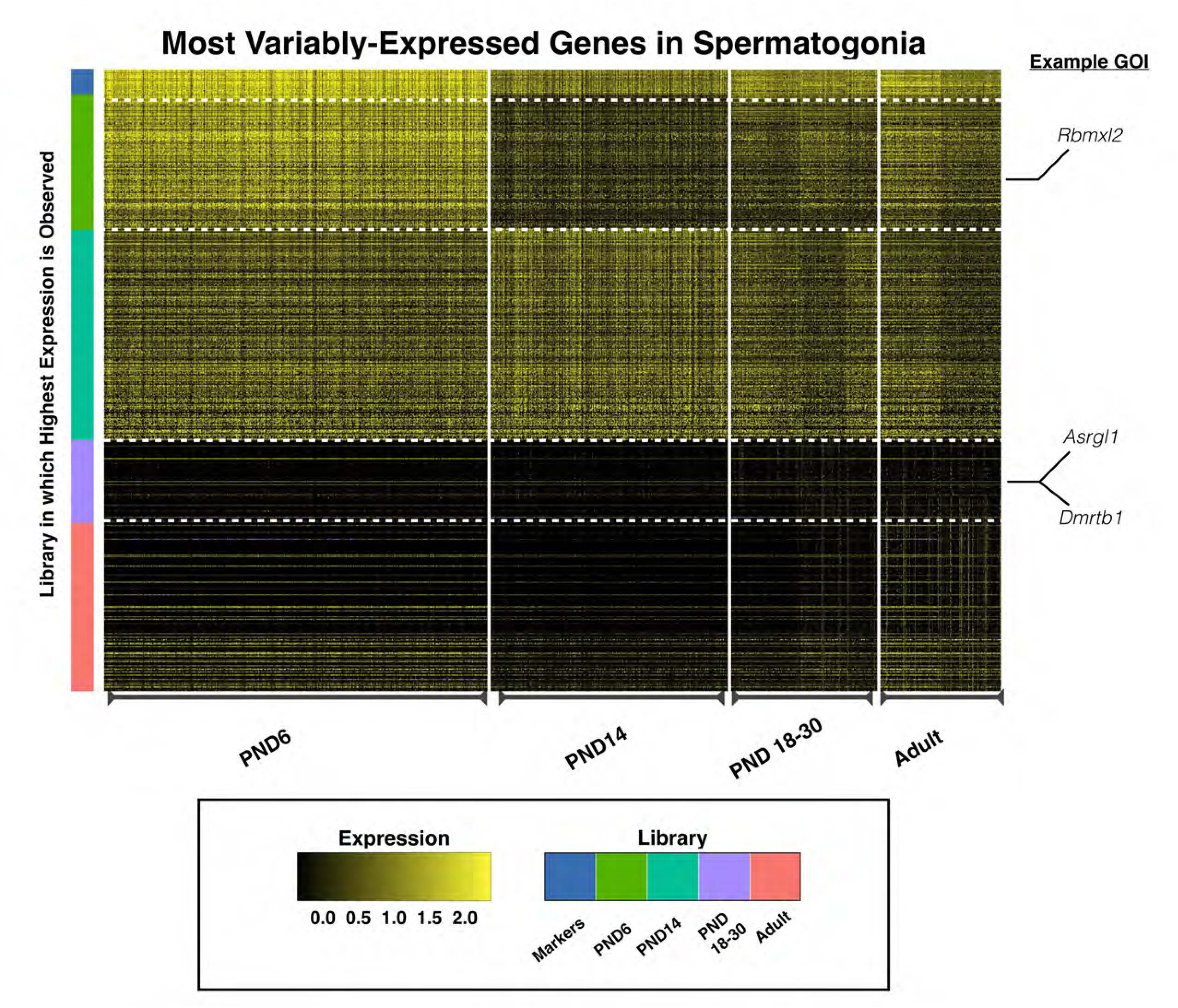
Analysis of variably-expressed genes in spermatogonia. MAST analysis was used to determine genes which are variably expressed with age specifically in spermatogonia, represented in a heatmap. All genes represented in the heatmap and listed in Table S3 are differentially expressed with the exception of the marker genes which remain consistently expressed. PND18-30 time points have been merged to increase representation of this rare cell type at those time points. Similarly, adult time points have also been merged. Individual cells are plotted vertically and the libraries from which they are derived is indicated at the bottom of the heatmap. Individual genes are plotted horizontally and the color bar at the left indicates library-of-origin from which highest expression is observed. Expression is scaled, ranging from 0 to 2.5.

In addition to MAST analysis for variable expression, Gene Set Enrichment Analysis (GSEA) Time Series analysis^20^ of Reactome pathways revealed differentially-utilized pathways, which were then visualized using Enrichment Map in Cytoscape^21,22^. GSEA of variably-expressed genes in spermatogonia from mice of different ages reveals significant changes in many pathways, including increasing expression of genes related to RNA destabilization and protein degradation as well as WNT signaling, and decreasing expression of genes related to asparagine metabolism, various signaling pathways including TGFB, FGFR, and KIT, and transcriptional regulation (**Figure 5 &S6A, Table S4**). In particular, many critical signaling receptors and ligands, including *Kit* and *KitL*^23–26^, as well as *Fgf8* and *Fgfr1*^27,28^, exhibit downregulation in spermatogonia derived from mice of increasing age, consistent with overall altered paracrine signaling around the basement membrane of the seminiferous tubules during testis maturation.

**Figure 5.**
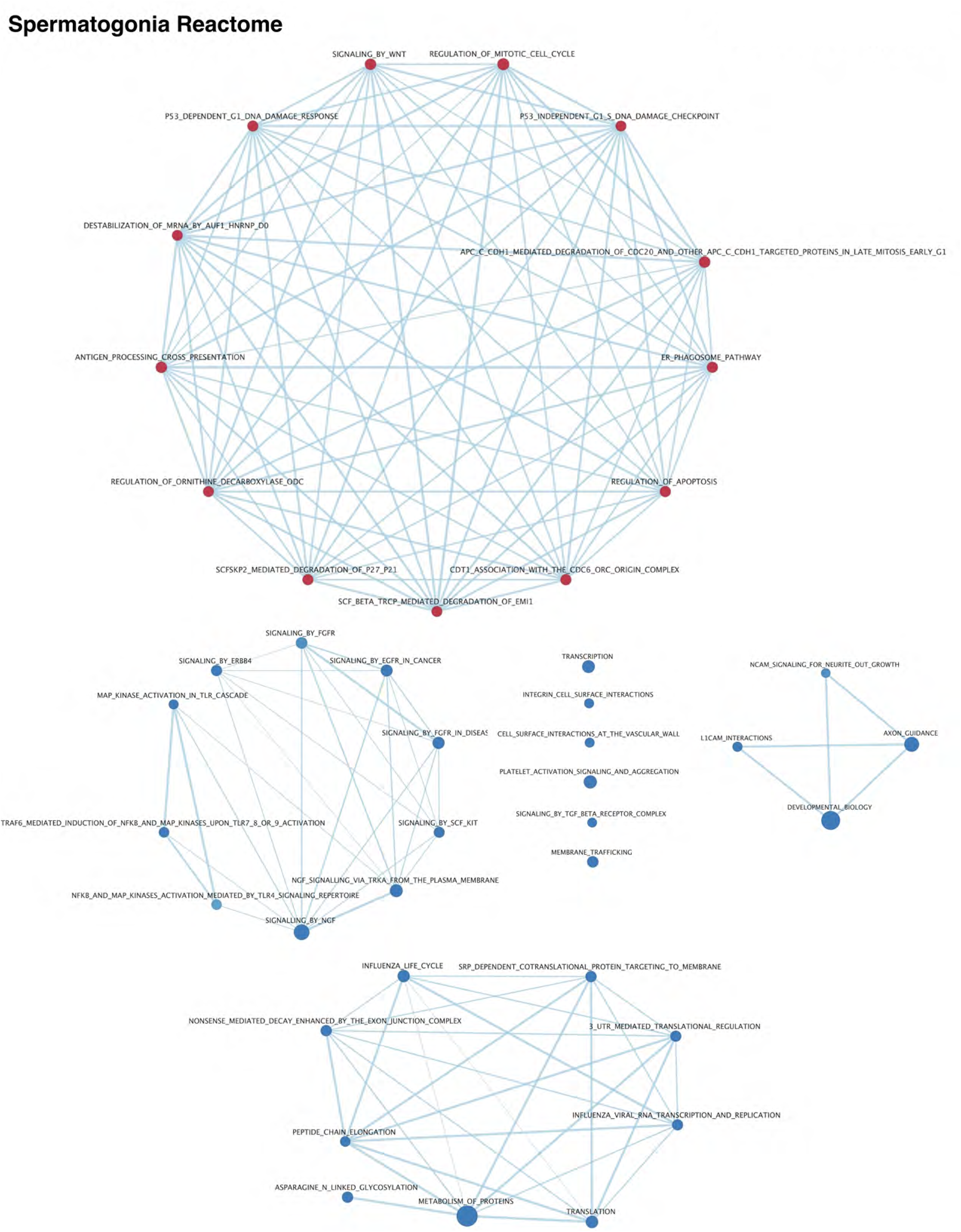
Differential Reactome pathway utilization in spermatogonia with age. Gene set enrichment analysis of variably-expressed genes in the Reactome database was visualized in Cytoscape. Results were filtered on a false discovery rate <0.05, and a gene set list >45 genes. Red nodes indicate pathways upregulated with time while blue nodes indicate pathways down-regulated with time. Edges indicate connections and overlap between pathways.

### Spermatocytes from the first wave of spermatogenesis are transcriptionally distinct from steady-state spermatocytes

It has been well established that meiotic regulation is distinct in the first wave of meiosis from that of subsequent waves^6,14,15^. Thus, we sought to explain this phenomenon in terms of the transcriptome profile of spermatocytes at discrete developmental time points. Spermatocytes from the first meiotic wave compared to steady-state (adult) ages were also subjected to MAST analysis, as described above (**Figure 6**, **Table S5**). For this analysis, spermatocytes were abundant enough from all libraries that each time point could be considered separately, except for PND6 in which spermatocytes are not yet present. Notably, spermatocytes from PND14, which are only just beginning Prophase I, demonstrate very distinct gene expression patterns from spermatocytes at later time points and are not representative of the full spectrum of meiotic cell types. Some genes, including those noted to the side of the heatmap, show robust differential expression with age, highlighting differences between spermatocytes derived from the first-wave (PND18) in contrast those which are derived from a self-renewing SSC population (adult). Therefore, these genes were chosen for further analysis and orthogonal validation.

**Figure 6.**
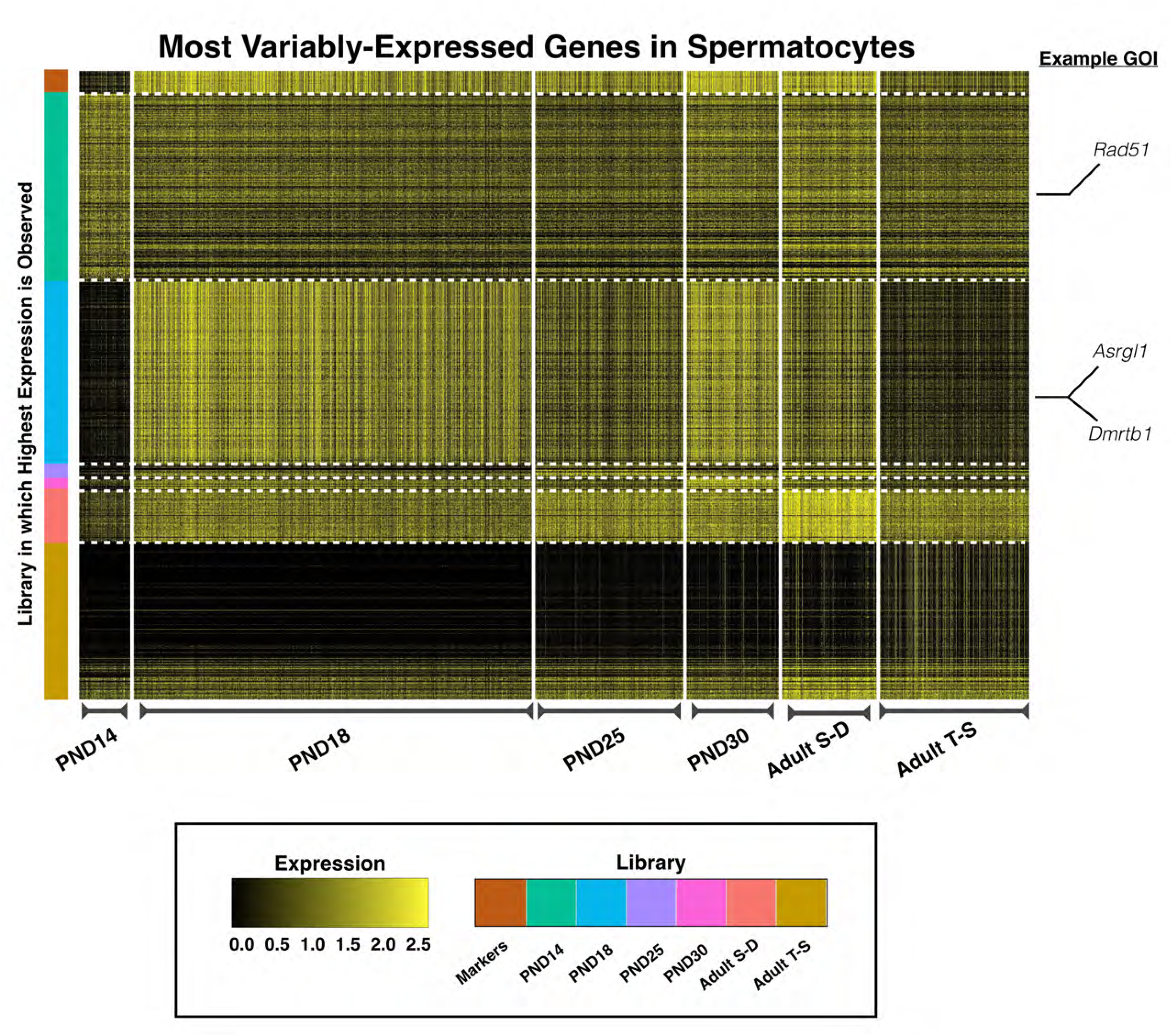
Analysis of variably-expressed genes in spermatocytes. MAST analysis was used to determine genes which are variably expressed with age specifically in spermatocytes, represented in a heatmap. All genes represented in the heatmap and listed in Table S5 are differentially expressed with the exception of the marker genes which remain consistently expressed. Individual cells are plotted vertically and the libraries from which they are derived is indicated at the bottom of the heatmap. Individual genes are plotted horizontally and the color bar at the left indicates library-of-origin from which the highest expression is observed. Expression is scaled, ranging from 0 to 2.5. The adult sperm-depleted sample is named “Ad S-D” while the adult THY1-sorted sample is named “Ad T-S”.

GSEA time series analysis of Reactome pathway enrichment of variably expressed genes in spermatocytes also reveals intriguing differentially utilized pathways. From this analysis, we observe decreasing expression of genes related to translation and post-transcriptional regulation, and increasing expression of genes related to DNA replication, double strand break repair, and cell cycle regulation (**Figure 7 & S6B, Table S6**). Most notable in the list of genes upregulated in spermatocytes of increasing age are those known to be essential to DNA repair, meiotic progression, and crossover formation including *Brip1*^29^, *Brca1* and *Brca2*^30–32^, *Rad51*^33^, *H2afx*^34^ and *Atm*^35^. Many of these pathways, particularly those related to double strand break repair (which initiates meiotic recombination), may be crucial for understanding the molecular mechanisms underlying fundamental differences in first-wave spermatocytes.

**Figure 7.**
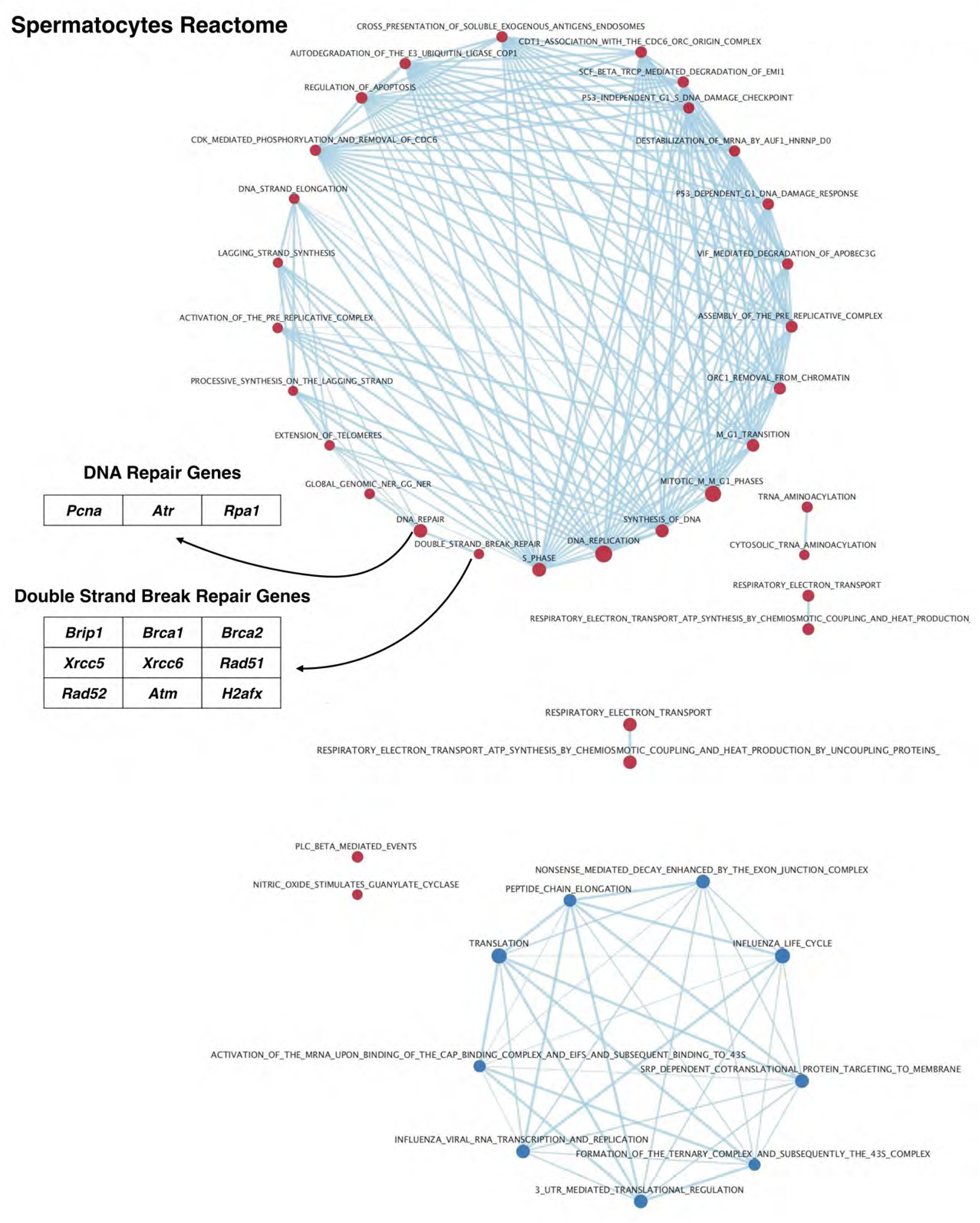
Differential Reactome pathway utilization in spermatocytes with age. Gene set enrichment analysis of variably-expressed genes in the Reactome database was visualized in Cytoscape. Results were filtered on a false discovery rate <0.05, and a gene set list >15 genes. Red nodes indicate pathways upregulated with time while blue nodes indicate pathways down-regulated with time. Edges indicate connections and overlap between pathways. The tables to the left of the diagrams identify notable genes represented in these pathways.

### Validation of differentially expressed genes demonstrates dynamic protein expression within a defined cell type during testis maturation

Candidate genes of interest (GOIs) identified in the single-cell sequencing data were investigated using immunofluorescence, which allowed us to validate changes in protein expression in the context of the native testis tissue. We used paraffin-embedded testis tissue sections at postnatal ages PND 7, PND 13, PND 22, and 8 weeks of age to characterize the same range of postnatal testis development as was captured in the single-cell RNAseq data set. GOIs were selected by meeting several criteria including: representation across several biological pathways, significantly differential expression in a given cell type between mice of different ages, and availability of a commercial immunofluorescence-verified and mouse-reactive antibody. For all validation of spermatogonial candidates, double-staining was performed with an antibody against ‘Promyelocytic Leukemia Zinc Finger Protein’ (PLZF; aka ZBTB16), a well-characterized marker of undifferentiated spermatogonia^36–38^. For all validation of spermatocyte candidates, double-staining was performed with an antibody against ‘Synaptonemal Complex Protein 3’ (SYCP3), to allow for visualization and staging of Prophase I-staged cells^13,39^. SYCP3 marks the nuclei of spermatocytes through leptotene, zygotene, pachytene, and diplotene stages of prophase I.

To profile a range of biological functions including metabolism, enzyme ‘Asparaginase-Like 1’ (ASRGL1; aka ALP1) was chosen for immunofluorescence analysis. ASRGL1 is known to catalyze the hydrolysis of L-asparagine^40^ and to clear protein-damaging isoaspartyl-peptides^41^, and while largely uncharacterized in the mouse, has been found to be highly expressed in the human cervix, fallopian tube, ovary, and testis^42^. Interestingly, ASRGL1 has been identified as a biomarker of endometrial cancer^42–45^, as well as an antigen in rodent sperm^46^. In the single-cell sequencing dataset, *Asrgl1* was observed to be highly expressed in spermatogonia from PND6 mice, with decreasing expression in this cell type of older mice (**Figure 4**). Interestingly, *Asrgl1* was also shown to have dynamic expression in spermatocytes, with the lowest expression detected in PND14 spermatocytes and increasing expression in spermatocytes of older mice (**Figure 6, Table S5**). These results at the mRNA level were corroborated by immunofluorescence data showing high expression of ASRGL1 protein in PND7 PLZF+ spermatogonia, with decreasing expression of the protein in PND22 and adult PLZF+ spermatogonia (**Figure 8A, S7**). Furthermore, PND13 first-wave pachytene spermatocytes express little ASRGL1 protein, with expression becoming abundant in pachytene spermatocytes from PND22 and adult testes (**Figure 8B, S8**).

**Figure 8.**
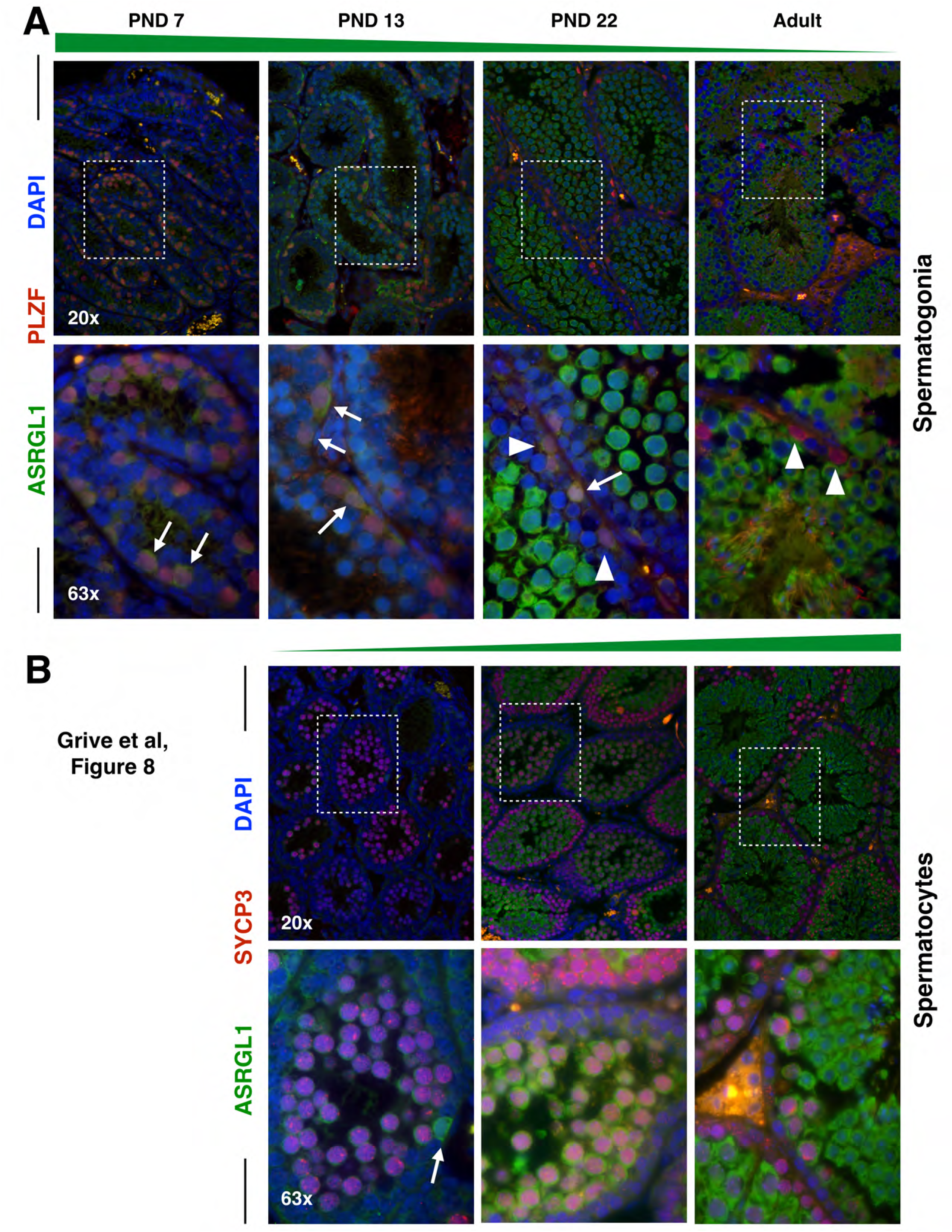
ASRGL1 is highly expressed specifically in spermatogonia from neonatal and juvenile mice, and spermatocytes of older mice. A) Spermatogonial marker PLZF (red) and ASRGL1 (green) were stained in 5μm testis tissue sections from mice ages PND7, PND13, PND22, and adult. DAPI (blue) denotes nuclei. ASRGL1 protein expression decreases in PLZF+ spermatogonia with age. B) Spermatocyte marker SYCP3 (red) and ASRGL1 (green) were stained in 5μm testis tissue sections from mice ages PND13, PND22, and adult. ASRGL1 protein expression increases in SYCP3+ spermatocytes with age. For all images, high-ASRGL1-expressing spermatogonia are indicated by full arrows with a line, while low-ASRGL1-expressing spermatogonia are indicated by arrowheads.

Significant differences in RNA stability and processing genes were also observed in spermatogonia during postnatal testis development, with down-regulation of related pathways over time. The RNA binding protein, ‘RNA Binding Motif Protein, X-linked-like 2’ (RBMXL2; aka HNRNPG-T) is a putative RNA regulator and splicing factor highly expressed in the mouse testis, specifically in germ cells^47^, with critical functions in spermatogenesis^48^. Furthermore, disruptions in RBMXL2 expression and localization in human testes are associated with azoospermia in men^49^. In the single-cell data set, *Rbmx2* mRNA was observed to be highly expressed in spermatogonia from the youngest mice, then decreasing with age, as well as expressed in all spermatocytes of all ages (**Figure 4, Table S3**). Immunofluorescence of RBMXL2 protein demonstrates high expression of the protein in all germ cells, including spermatogonia and spermatocytes. Close inspection, however, reveals relatively higher expression of RBMXL2 in PND7 PLZF+ spermatogonia compared to later time points, despite the relatively consistent expression of RBMXL2 protein in all other germ cell stages at all mouse ages (**Figure S9**).

‘Double-sex and Mab-3 Related Transcription Factor B1’ (DMRTB1; aka DMRT6) is a transcriptional regulator known to coordinate the developmental transition from spermatogonial differentiation to meiotic entry^50^. As has been previously observed, *Dmrtb1* mRNA is highly expressed in spermatogonia and early spermatocytes (**Figures 4 & 6**), which we have confirmed at the protein level by immunofluorescence. At PND13, first-wave early leptotene spermatocytes, evidenced by spotty SYCP3, exhibit nuclear DMRTB1 staining, while pachytene spermatocytes from all mouse ages lose DMRTB1 expression. Interestingly, the nuclear staining in leptotene spermatocytes is only observed at the earliest time point, PND13, and not seen in early Prophase I spermatocytes of later spermatogenic waves (**Figure 9, S10**).

**Figure 9.**
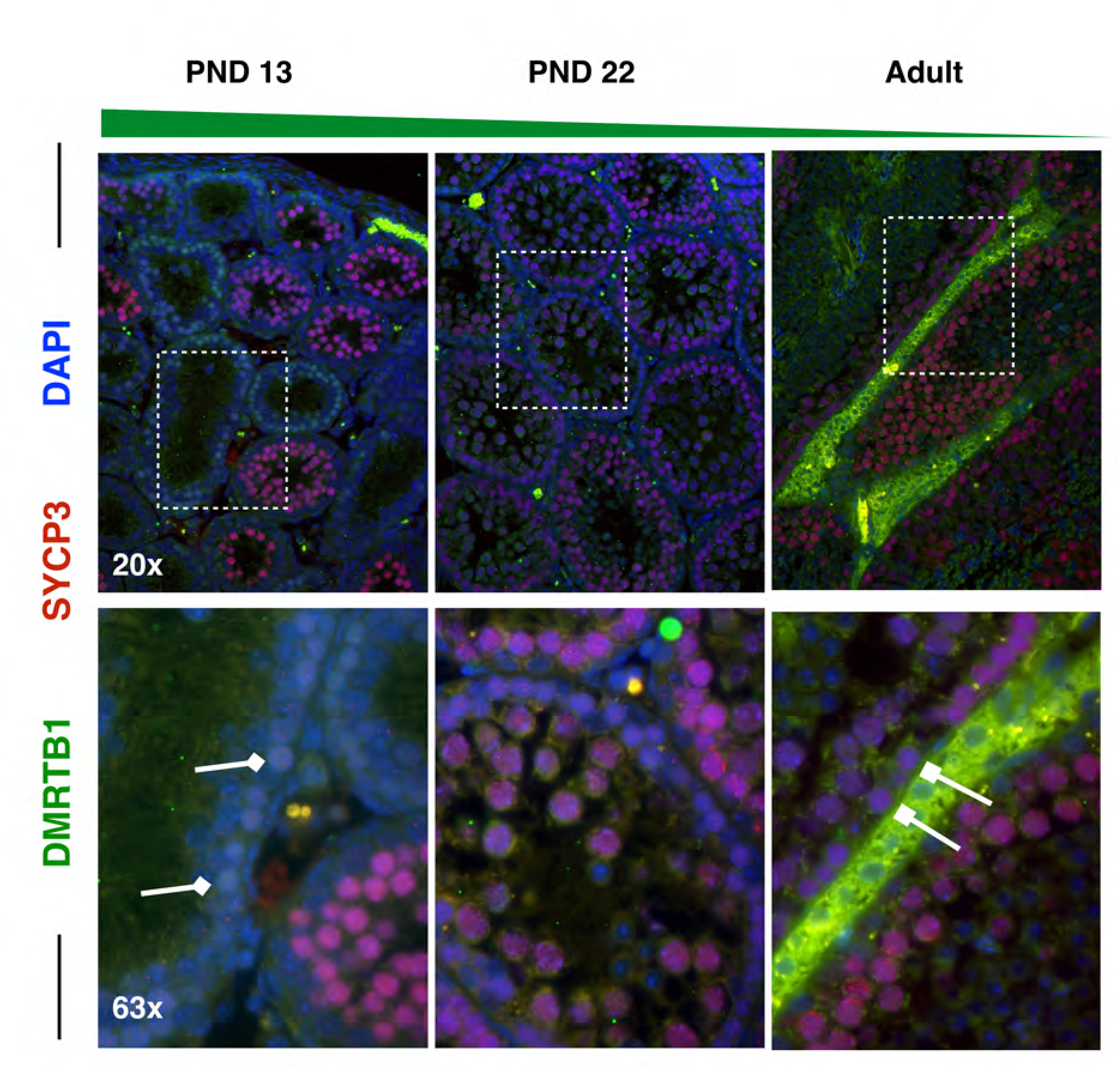
DMRTB1 is highly expressed specifically in first-wave spermatocytes from juvenile mice. Spermatocyte marker SYCP3 (red) and DMRTB1 (green) were stained in 5μm testis tissue sections from mice ages PND13, PND22, and adult. DMRTB1 protein is expressed in the nucleus of first-wave spermatocytes at PND13, with decreasing expression in pachytene spermatocytes with age. For all images, high-DMRTB1-expressing spermatocytes are indicated by diamond-headed arrows, while low-DMRTB1-expressing spermatocytes are indicated by square-headed arrows.

Finally, DNA damage response proteins ‘RAD51 Recombinase’ (RAD51) and ‘Ataxia Telangiectasia Mutated’ (ATM) were profiled across spermatocytes from mice of increasing age, as these represent particularly interesting candidate proteins whose differential expression may be crucial to understanding aberrant recombination rates and chromosome segregation in the first wave. Intriguingly, both RAD51 and ATM show subtly, but decidedly, decreased expression in first-wave pachytene spermatocytes, with increasing expression as mice age (**Figures 10, S11, S12**), as predicted from our scRNAseq data. These observations of altered RAD51 staining intensity are consistent with previous reports of significantly fewer RAD51 foci along chromosome cores of juvenile C57bl/6 spermatocytes compared to those at 12 weeks of age^14^. Our RAD51 observations demonstrate overall decreased protein expression in the nucleus of pachytene first-wave spermatocytes, with increasing expression by 3 weeks of age (**Figure 10B, S12**). A similar dynamic is observed for ATM, which has robust cytoplasmic staining as well as diffuse nuclear staining in pachytene spermatocytes^51^. Our data demonstrate decreased expression of ATM in both cellular compartments in the first-wave spermatocytes, with increasing expression, particularly in the cytoplasm of these cells, by 3 weeks of age (**Figure 10A, S11**). These age-dependent dynamics of critical DNA damage response regulators are likely to contribute to the health and viability of resulting spermatocytes and spermatozoa from these spermatogenic cycles, and may underlie some of the functional differences observed in the first wave of spermatogenesis.

**Figure 10.**
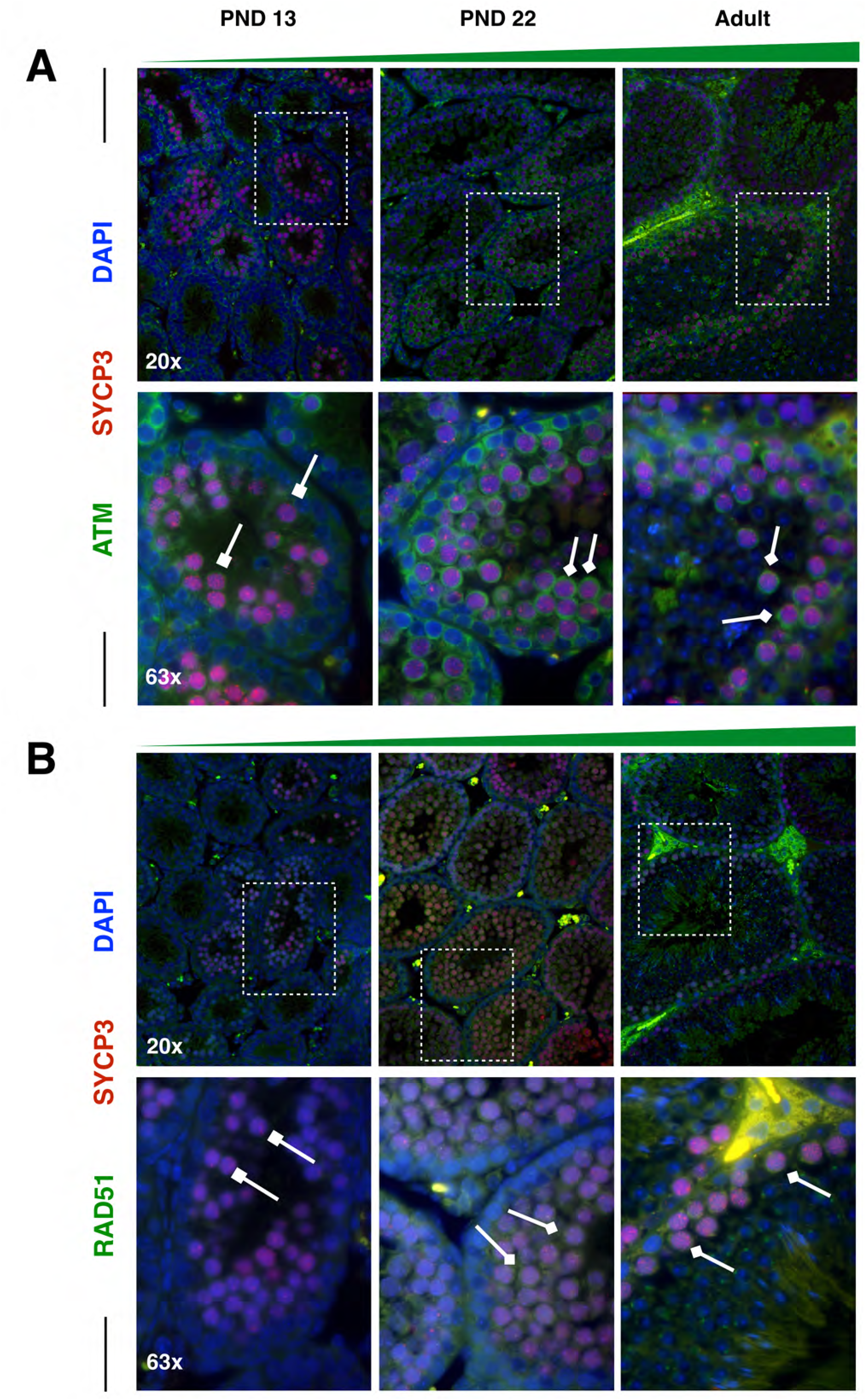
ATM and RAD51 have reduced expression in first-wave spermatocytes from juvenile mice. A) Spermatocyte marker SYCP3 (red) and ATM (green) were stained in 5μm testis tissue sections from mice ages PND13, PND22, and adult. ATM protein expression increases in SYCP3+ spermatocytes with age. For all images, high-ATM-expressing spermatocytes are indicated by diamond-headed arrows, while low-ATM-expressing spermatocytes are indicated by square-headed arrows. B) Spermatocyte marker SYCP3 (red) and RAD51 (green) were stained in 5μm testis tissue sections from mice ages PND13, PND22, and adult. RAD51 protein expression increases in SYCP3+ spermatocytes with age. For all images, high-RAD51-expressing spermatocytes are indicated by diamond-headed arrows, while low-RAD51-expressing spermatocytes are indicated by square-headed arrows.

Overall, these gene expression dynamics discovered from single-cell mRNA sequencing are reproducible at the level of protein expression in the context of the native tissue, and likely represent important development transitions both in spermatogonia and spermatocytes. These data will be indispensable to investigate how gene expression dynamics help to regulate the many critical developmental events, including spermatogonial differentiation and meiotic progression, occurring in the developing mouse testis.

## Discussion

We have performed the first comprehensive sampling and screening of mouse germ cells from neonatal life through adulthood, to obtain transcriptional profiles at a high, single-cell resolution. With the exception of the adult samples, which were sorted to exclude the over-riding sperm component, all libraries were generated from single-cell suspensions of all testicular cells. Adult samples were minimally processed with a single-step magnetic cell sort to provide representation of all germ cell types in the adult testis. Previously, single-cell-sequencing studies on sorted cells have provided valuable information about specific, and marker-defined cell types. The study presented here, however, focuses on profiling the germline during postnatal testis maturation, importantly capturing the first-wave of spermatogonia and spermatocytes which exhibit differences from later, steady-state spermatogenesis. Because of the progression of ages profiled, we have captured changes in gene expression at single-cell resolution to compare the developmental progression of spermatogenesis as mice age.

Germ cells subtypes in the testis are frequently defined by the presence or absence of particular protein markers, which can be visualized by reporter expression or immunofluorescence, or which can sometimes be used for flow cytometry or other enrichment paradigms. Spermatogonia are often defined as cells which express a key complement of protein markers, such as PLZF or ‘GDNF family receptor alpha 1’ (GFRα1)^52–54^. While this is the best practice for visual identification of cells for which discrete markers have been elusive, such as spermatogonia, our analysis suggests that the biology of these cells during postnatal testis development is far more complicated that previously understood. Our analysis stresses that defining these cell populations on the basis of specific markers may be overly simplistic, despite being the current standard practice in the field. Primary spermatocytes are similarly distinct at the transcriptional level over developmental time. Furthermore, cells possessing an SYCP3-positive synaptonemal complex indicative of pachynema also exhibit differences in immunolocalized proteins during testis maturation, indicating that they, too, exhibit distinct and variable translational dynamics with increasing age. Critically, this analysis reveals that, while known markers may be useful for defining primary cell identity, there are many changes in spermatogenesis that have been under-appreciated without the power of single-cell resolution of gene expression profiling.

Specifically, we show here that PLZF-defined spermatogonia, though retaining PLZF-positivity, are transcriptionally distinct at PND6 compared to later developmental time points. These transcriptional dynamics are also reflected by distinct differences at the protein level, with proteins such ASRGL1 being localized strongly in PLZF+ spermatogonia during the first weeks of life, but decreasing in expression in these cells around three weeks of age (**Figure 8A, S7**). Similarly, we show that Prophase I spermatocytes possess significant transcriptional differences in the first-wave compared to subsequent spermatogenic waves, with hundreds of differentially expressed genes across multiple regulatory pathways. Furthermore, direct inspection of pachytene spermatocytes from the first wave at PND13 to later spermatogenic waves reveals that, like spermatogonia, transcript dynamics are also reflected at the protein level. For example, proteins such as DMRTB1 are found only in the nuclei of first-wave leptotene spermatocytes as they transition from differentiated spermatogonia into the meiotic program, but not in spermatocytes from older mice (**Figure 9, S10**). By contrast, other proteins such as ASRGL1 are in low abundance in early first-wave spermatocytes, but become more strongly immunolocalized in spermatocytes at increasing ages (**Figure 8B, S8**).

In addition to the transcriptional and translational dynamics in defined cell types over time, these data also reveal differential utilization of particular biological pathways over developmental time. Gene set enrichment analysis^20^ utilizing the Reactome pathway database^21,22^ has demonstrated that spermatogonia dramatically change their transcriptional landscape as mice age, including downregulation of genes within essential meiotic-entry-associated SCF/KIT^26,55^ and FGFR^56^ pathways, including *Kit* and *KitL*. While we cannot rule out the possibility that variable gene expression in spermatogonia is, in part, due to differential contributions of the spermatogonial stem cell population at different ages – with decreasing contribution as mice age – this dataset provides strong support for true variable gene expression in the spermatogonial pool. For instance, while spermatogonia derived from older mice exhibit downregulation of genes associated with FGFR signaling, including *Fgf8* and *Fgfr1*, and could indicate decreased representation of an undifferentiated spermatogonial population^28^, this is in opposition to observed coincident decreased expression of *Kit* and *KitL* which would support increased representation of an undifferentiated population^24,26,55^. Therefore, these data likely reflect overall changes to the paracrine signaling of the spermatogonial stem cell niche as well as the basement membrane as mice age. Overall, these data suggest spermatogonia may modulate their sensitivity for particular critical signaling pathways, which may affect their competency to commit to the meiotic program. Furthermore, pathways associated with mRNA stability and protein degradation are upregulated as the testis matures, suggesting that spermatogonia from older mice may change their capacity for post-transcriptional and post-translational regulation with age, possibly reflecting changing demands for growth and proliferation in older animals.

Similar to spermatogonia, spermatocytes also exhibit differential utilization of specific biological pathways with age, an observation that dovetails with the knowledge that spermatocytes derived from the first wave of spermatogenesis are functionally different to those spermatocytes derived from steady-state spermatogenesis. First-wave spermatocytes are known to exhibit several unique, and some detrimental, characteristics, including reduced recombination rate^14,15^ and greater incidence of chromosome mis-segregation^15^. These features result in first-wave spermatozoa which are often much less reproductively successful than those which will arise from the self-renewing SSC population later in life^57,58^. Our data presented here demonstrate age-related upregulation of pathways associated with DNA replication and repair, double strand break repair, and response to DNA damage, all of which may underlie the well-characterized differences between spermatocytes in the first wave compared to steady-state spermatocytes. Included in these sets of variably-expressed genes are known regulators of DNA damage response and cross-over formation including *Rad51*^33^*, Brip1*^29^*, and H2afx*^34^, as well as *Brca1* and *Brca2*^30–32^ and *Atm*^35,59^, all of which increase in spermatocytes with age, effects which we have also shown at the protein level for both RAD51 and ATM (**Figures 10, S11, S12**). While the cause of this lower expression in the first wave is unknown, it has been demonstrated that spermatocytes from juvenile mice generate only about 25% of the double strand breaks present in spermatocytes from steady-state spermatogenesis^59^. It is possible that in response to fewer breaks, less DNA damage response machinery is required and therefore exhibits lower expression levels. Alternatively, these observations may also suggest that steady-state spermatocytes acquire greater competency to cope with the DNA damage inherent to meiotic progression, and that spermatocytes in the first wave may not execute these pathways as successfully, resulting in the observed recombination differences and increased chromosome mis-segregation. Notably, like spermatogonia, spermatocytes also experience alteration of pathways related to translation and mRNA stability, emphasizing the myriad ways in which gene expression is regulated in developing germ cells. Ultimately, this differential pathway utilization may help to explain not only the functional differences observed in spermatocytes and spermatozoa from juveniles, but may also improve our understanding of increased birth defects associated with young paternal age^57,58^.

Another of our primary objectives in undertaking this analysis was to potentially reveal new markers of the spermatogonial stem cell population. Despite many efforts to define this population by both cytoplasmic and nuclear markers, discrete markers of this population have remained elusive and controversial ^60–63^. Our data are supportive of the high degree of heterogeneity of this cell population, not only within the population at a single age, but also across ages. Furthermore, markers which have become accepted in the field, such as expression of ‘Inhibitor of DNA Binding 4’ (*Id4*)^64–66^, show widespread detection in spermatogonia through spermatocytes, while markers such as *Zbtb16* (*Plzf*) and *Gfr*α*1* have much more restricted expression (**Figure S5**). Importantly, expression patterns of these popular markers are not entirely self-consistent. It is therefore likely that the spermatogonial population is not discrete, but is indeed a continuous or plastic population^67,68^. Despite this, these data remain a valuable resource to those interested in understanding the molecular mechanisms underlying SSC self-renewal and differentiation, though the biology may not be as simplistic as originally thought.

Taken together, these data represent the first comprehensive sampling and profiling of spermatogonia and spermatocytes during development of the mouse testis. These data emphasize the necessity of considering not only the protein markers for which individual cells are positive, but also the age of the cells being analyzed. These observations of highly dynamic gene expression in germ cell populations during postnatal testis development stress that germ cells of a particular age or identity possess vastly different underlying biology and that consideration must be given to these dynamics when profiling germ cell populations. These data also represent an invaluable community resource for discovery of previously unknown gene expression dynamics and pathway contributions that may be critical for the many developmental transitions in the male germ cell population which are essential for successful spermatogenesis and fertility.

## Methods

### Animals

B6D2F1/J mice were generated by mating C57bl/6 female mice with DBA/2J male mice. All animal protocols were reviewed and approved by the Cornell University Institutional Animal Care and Use Committee and were performed in accordance with the National Institutes of Health Guide for the Care and Use of Laboratory Animals. Mice were maintained on standard light:dark cycles with laboratory mouse chow provided *ad libidum.*

### Generation of testis single cell libraries

Testes were collected from mice (n = 1 mouse, 2 testes for each time point) at postnatal (PND) days 6, 14, 18, 25, 30, and 8 weeks of age, and dissociated per standard protocols for germ cell enrichment^16^. Briefly, testes were held in 1X HBSS buffer before de-tunicating and moving tubules into 0.25% Trypsin. Tubules were further dissociated by trituration and addition of DNase to a final concentration of 7 mg/ml. Tubules were placed in a 37°C incubation for 5 minutes at a time, and then removed for further trituration. Incubations at 37°C were performed three times, until a cloudy suspension was achieved. Cells were passed through a 40 μM filter, spun down, and re-suspended in 10ml 1X Dulbecco’s PBS + 10% Knockout Serum Replacement (DPBS-S). This cell suspension was then layered on top of a 30% Percoll solution. Cells were then spun down again, and the resulting pellet was re-suspended in 1ml DPBS-S. As a technical control, cells from PND18 were split into two samples after the 40 μM filter, with one half of the cells processed with the Percoll gradient, and the other half directly re-suspended in its final buffer with no Percoll sedimentation, resulting in libraries “PND18” and “PND18pre”, respectively. Due to the similarities between these libraries (**Figure S1**), the data from these libraries were thereafter combined and analyzed together as “PND18”.

For adult testes only, the resulting cell suspension was split in half and sorted with magnetic beads in two ways: (1) sperm-depletion was performed by incubating the cells for 30 minutes with 20 μl anti-ACRV1-PE (Novus Biologicals #NB500-455PE), washing with DPBS-S, incubating the cells for 30 minutes with 20 μl magnetic-bead-conjugated anti-PE (Miltenyi Biotec #130-048-801), and finally performing a negative magnetic selection. Cells were applied to a Miltenyi Biotec MACS LS column, and flow-through cells were collected, as sperm were to remain bound to the ferromagnetic column. (2) THY1+ spermatogonia were enriched by incubating the cells for 60 minutes with 20 μl magnetic-bead-conjugated anti-CD90.2 (THY1) (Miltenyi Biotec #130-102-464), and finally performing a positive magnetic selection. Cells were applied to the column, flow-through cells were discarded, and antibody-bound cells were eluted from the ferromagnetic column. These cells were then spun down and re-suspended in 1ml DPBS-S as above.

The resulting cells from all samples were submitted to the Cornell DNA Sequencing Core Facility for processing on the 10X Genomics Chromium System with a target of 4-5000 cells per sample. Sequencing libraries were generated using the 10X Genomics Chromium Single Cell 3′ RNAseq v2 kit, tested for quality control on an ABI DNA Fragment Analyzer, and run on a NextSeq platform with 150 base-pair reads. Libraries were sequenced to average depth 98M reads (range 77M-124M); on average, 91% of reads (range 89%-92%) mapped to the reference genome.

### Single-cell transcriptome analysis

Data normalization, unsupervised cell clustering, and differential expression were carried out using the Seurat R package^69^. Batch effect and cell-cycle effect were removed by Combat^70^ and Seurat together. Cells with less than 500 genes or 2000 UMIs or more than 15% of mitochondria genes were excluded from the analysis. Gene expression raw counts were normalized following a global-scaling normalization method with a scale factor of 10,000 and a log2 transformation, using the Seurat NormalizeData function. The top 4000 highly variable genes were selected using the expression and dispersion (variance/mean) of genes. Combat removed batch effects. Seurat regressed the difference between the G2M and S phase, then followed by principal component analysis (PCA). The most significant principal components (1-30) were used for unsupervised clustering and t-Distributed Stochastic Neighbor Embedding (tSNE) analysis.

Cell types were manually identified by marker genes^71–73^, and confirmed by SingleR (Single-cell Recognition) package. Differential expression analysis was performed based on the MAST (Model-based Analysis of Single Cell Transcriptomics)^19^. Gene Set Enrichment Time Series Analysis^20^ used the differential expression based on each time point, after removing genes highly expressed in spermatids. Pathways were visualized by EnrichmentMap^22^ in Cytoscape^21^.

#### Code availability

The scripts used for analysis and figure generation are available at https://github.com/nyuhuyang/scRNAseq-SSCs

#### Data availability

The single-cell RNAseq data have been deposited at GEO and are accessible through Series accession number: GSE121904.

### Immunofluorescence validation

Testes were collected, cleaned of excess fat, and fixed in 0.1% formalin solution overnight before dehydration and embedding in paraffin. Fixed testes were sectioned at 5 μm onto glass slides by the Cornell Animal Health Diagnostic Center. To stain, sections were de-paraffinized by 3x, 5 minute washes in Histoclear followed by rehydration in 100% ethanol (2x, 5 minutes), 95% ethanol (2x, 5 minutes), 70% ethanol (1x, 5 minutes), water (1x, 5minutes). Sections were then incubated in boiling antigen retrieval buffer (10 mM sodium citrate, 0.05% Tween-20, pH 6.0) for 20 minutes and left to cool. Sections were washed 3x, 5 minutes in 1X PBS + 0.1% Triton-X (PBST). Tissue sections were then incubated in blocking buffer [3% Goat Serum (Sigma), 1% Bovine Serum Albumin (Sigma), and 0.5% Triton-X (Fisher Scientific) in 1X PBS] and stained by incubation with primary antibodies against PLZF, SYCP3, RBMXL2, ASRGL1, DMRTB1, RAD51, and ATM (see **Table S7**) overnight at 4°C. The following day, slides were washed 3x, 5 minutes in PBST and then incubated with secondary antibodies raised in goat against mouse (594 nm) and rabbit (488 nm) at 1:500 for 1 hour at 37°C. A secondary antibody-only control was included to assess background staining. Sections were further stained with DAPI to visualize nuclei, mounted and analyzed on an Epifluorescent Zeiss Axioplan microscope. For all time points for a given set of antibodies, images were exposed equivalently.

---

**Supplementary Figure 1.**
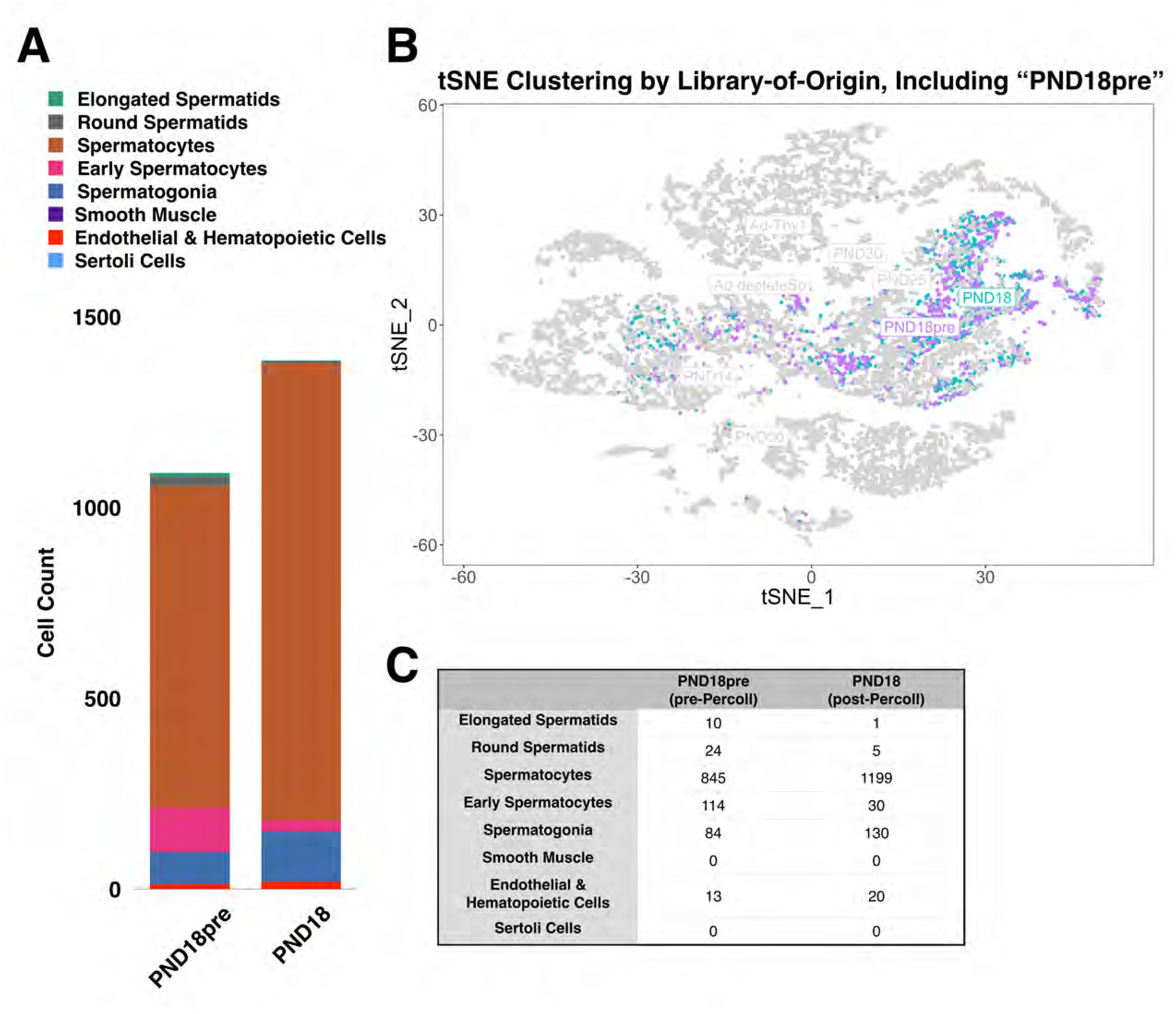
Similarities between “PND18pre” and “PND18” libraries. A) Germ and somatic cell composition by proportion and absolute cell number from libraries “PND18pre” (pre-Percoll) and “PND18” (post-Percoll). B) tSNE representation of all cells with >500 detected genes and >2000 UMIs. PND18 libraries are color-coded while other libraries are greyed out. C) Cell counts for each cell type plotted in (A). As a result of all of these similarities, the data derived from the libraries was combined for analysis.

**Supplementary Figure 2.**
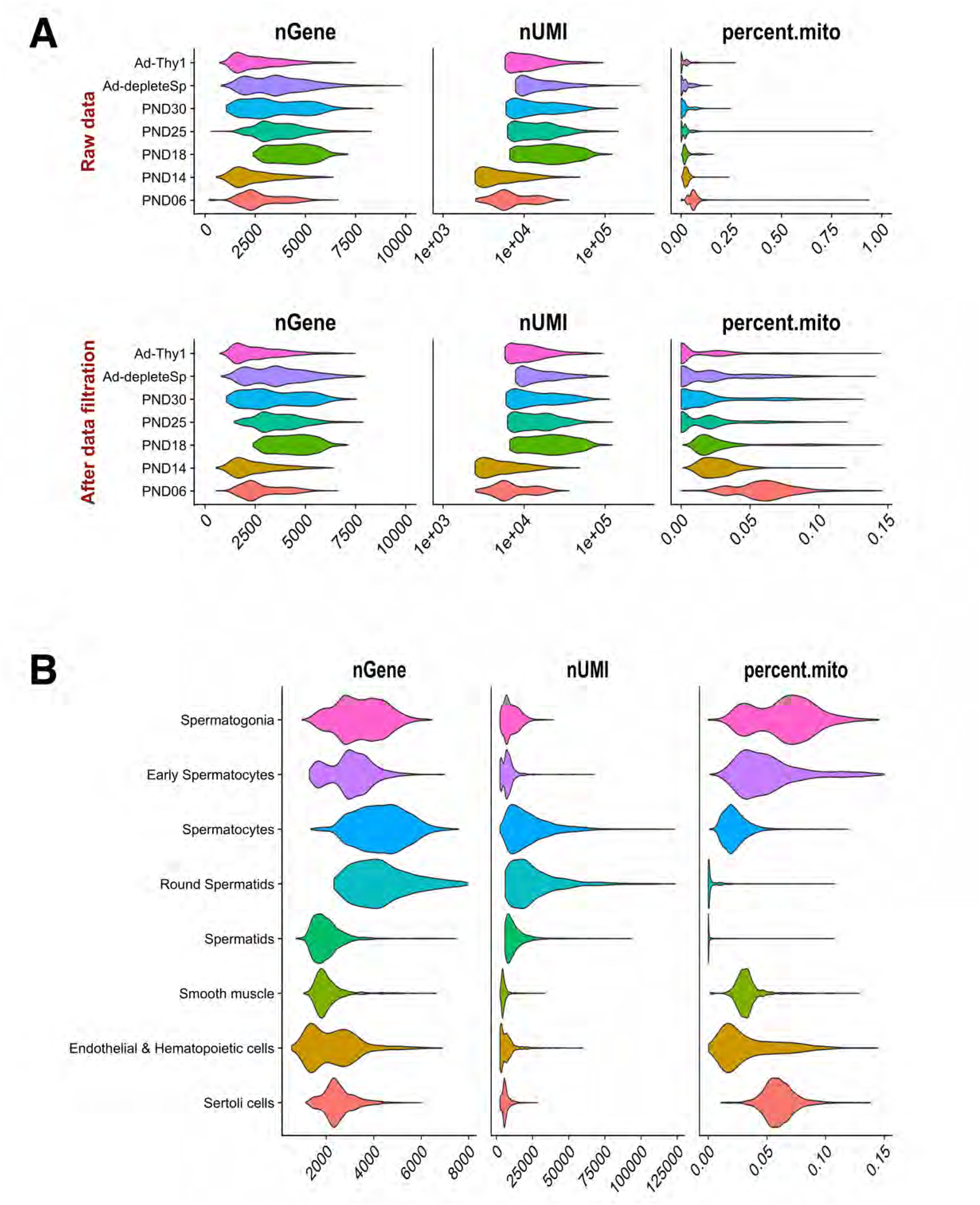
Quality control violin plots of single-cell data before and after filtering. A) Distribution of gene and UMI counts, and mitochondrial gene percentage per library-of-origin, before and after data filtration. B) Distribution of gene and UMI counts, and mitochondrial gene percentage per cell type.

**Supplementary Figure 3.**
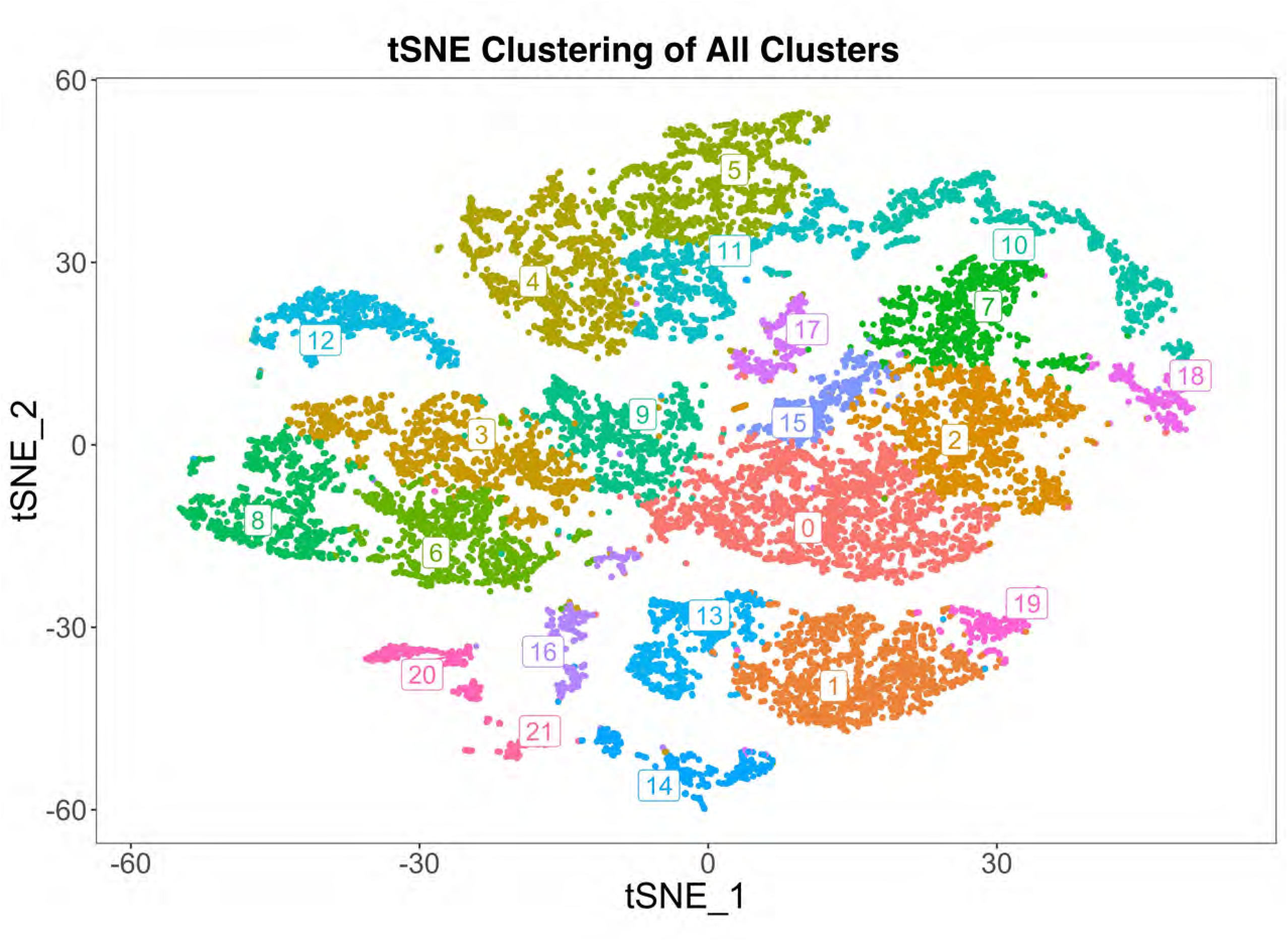
Additional clustering of data, into computationally determined clusters. Representative clustering of all cells with >500 detected genes and >2000 UMIs, based on most significant principal components, color-coded by cell cluster.

**Supplementary Figure 4.**
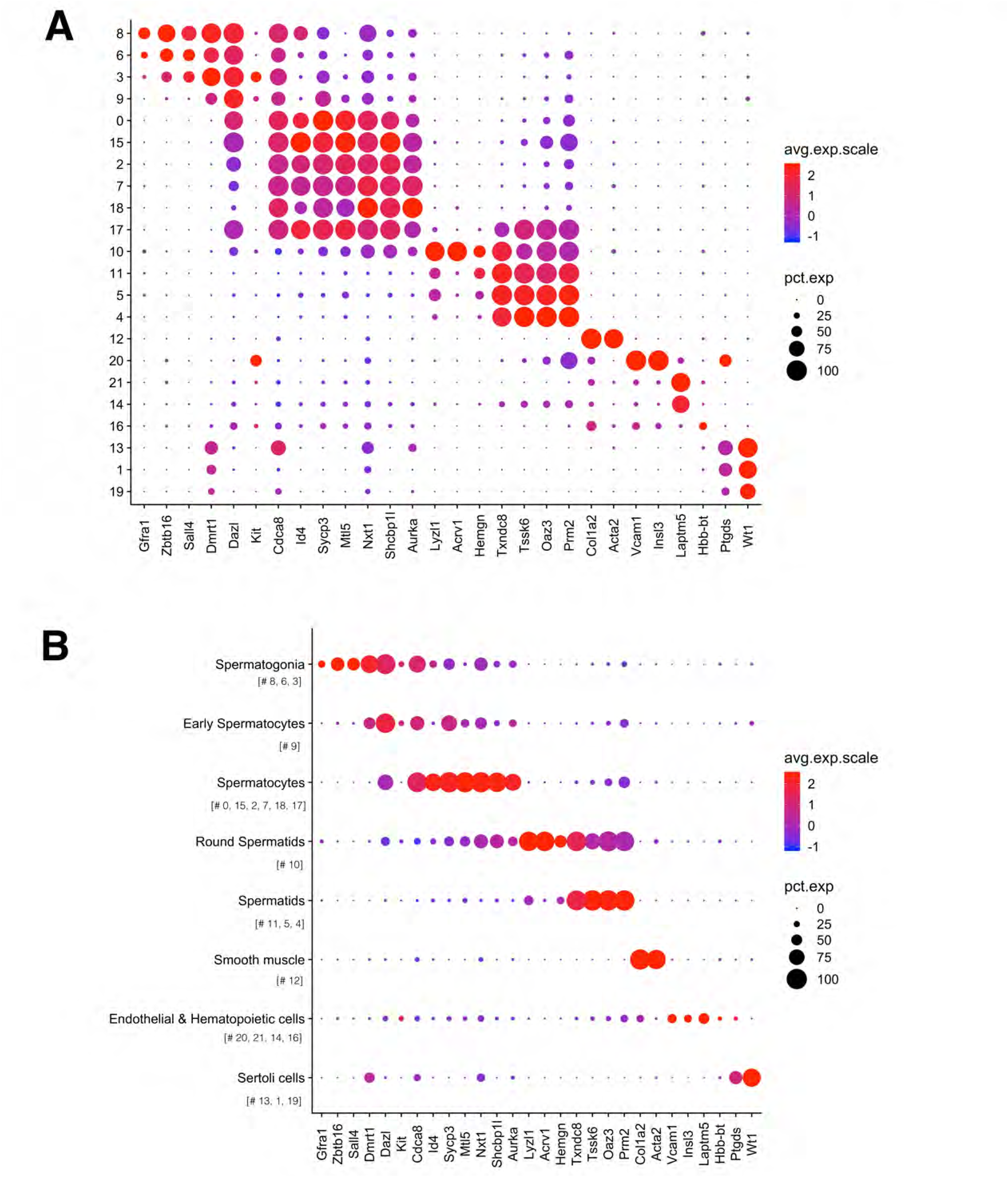
Dot plot analysis of known gene expression markers based on cluster and cell type. A) Dot plot representation of known marker genes per cell cluster determined in Figure S3. B) Dot plot representation of known marker genes per cell type determined in Figure 2.

**Supplementary Figure 5.**
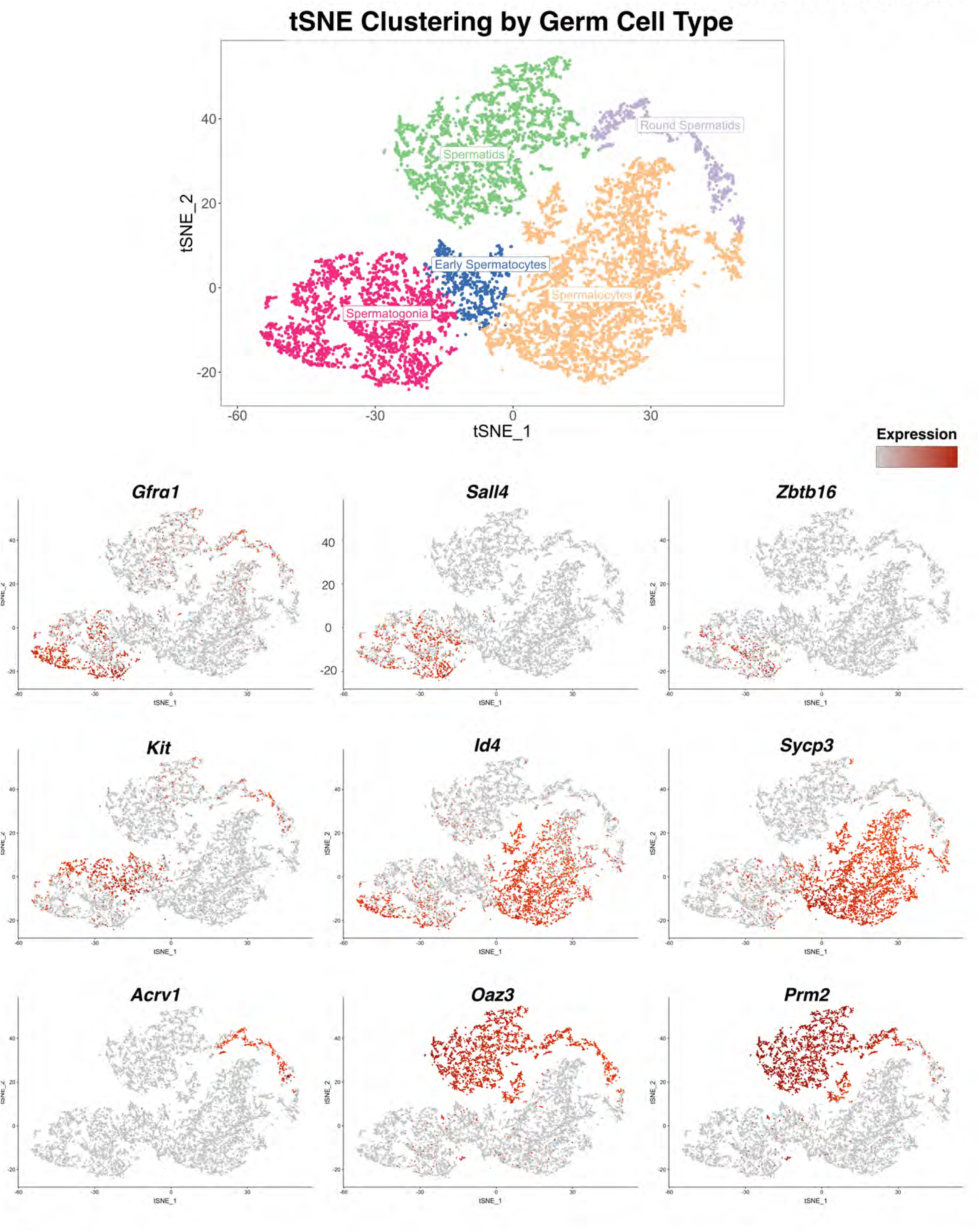
Representative germ cell marker expression in cells from all libraries-of-origin. tSNE plot of germ cells from all libraries, as well as tSNE plots of notable germ cell marker gene expression.

**Supplementary Figure 6.**
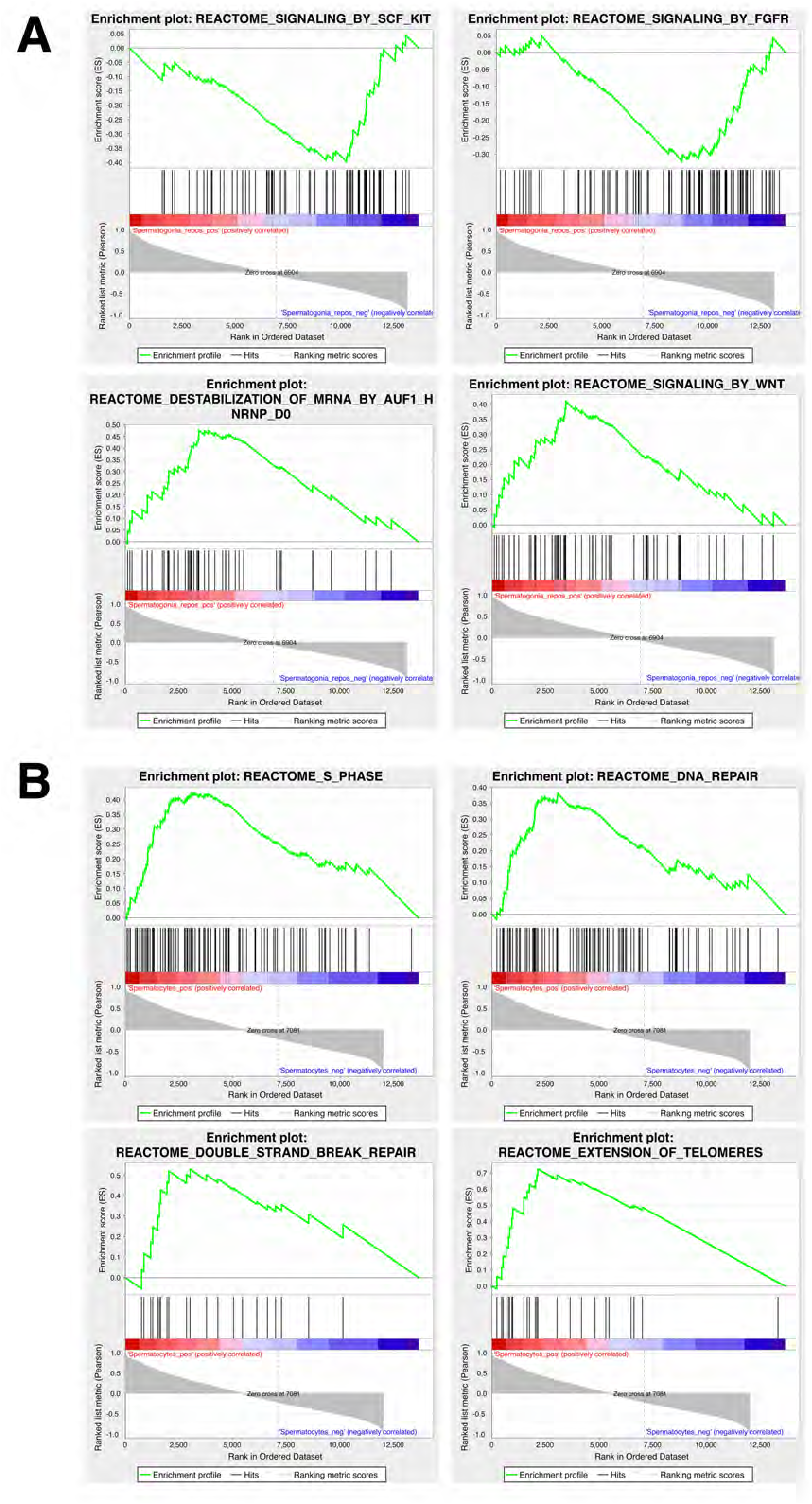
GSEA enrichment plots for pathways correlated with spermatogonia and spermatocyte development. A) Enrichment plots for selected Reactome database pathways in spermatogonia. Pathways “SIGNALING_BY_SCF_KIT” and “SIGNALING_BY_FGFR” show negative correlation with developmental time, while pathways “DESTABILIZATION_OF_MRNA” and “SIGNALING_BY_WNT” show positive correlation with developmental time. B) Enrichment plots for selected Reactome database pathways in spermatogonia. All pathways shown demonstrate positive correlation with developmental time.

**Supplementary Figure 7.**
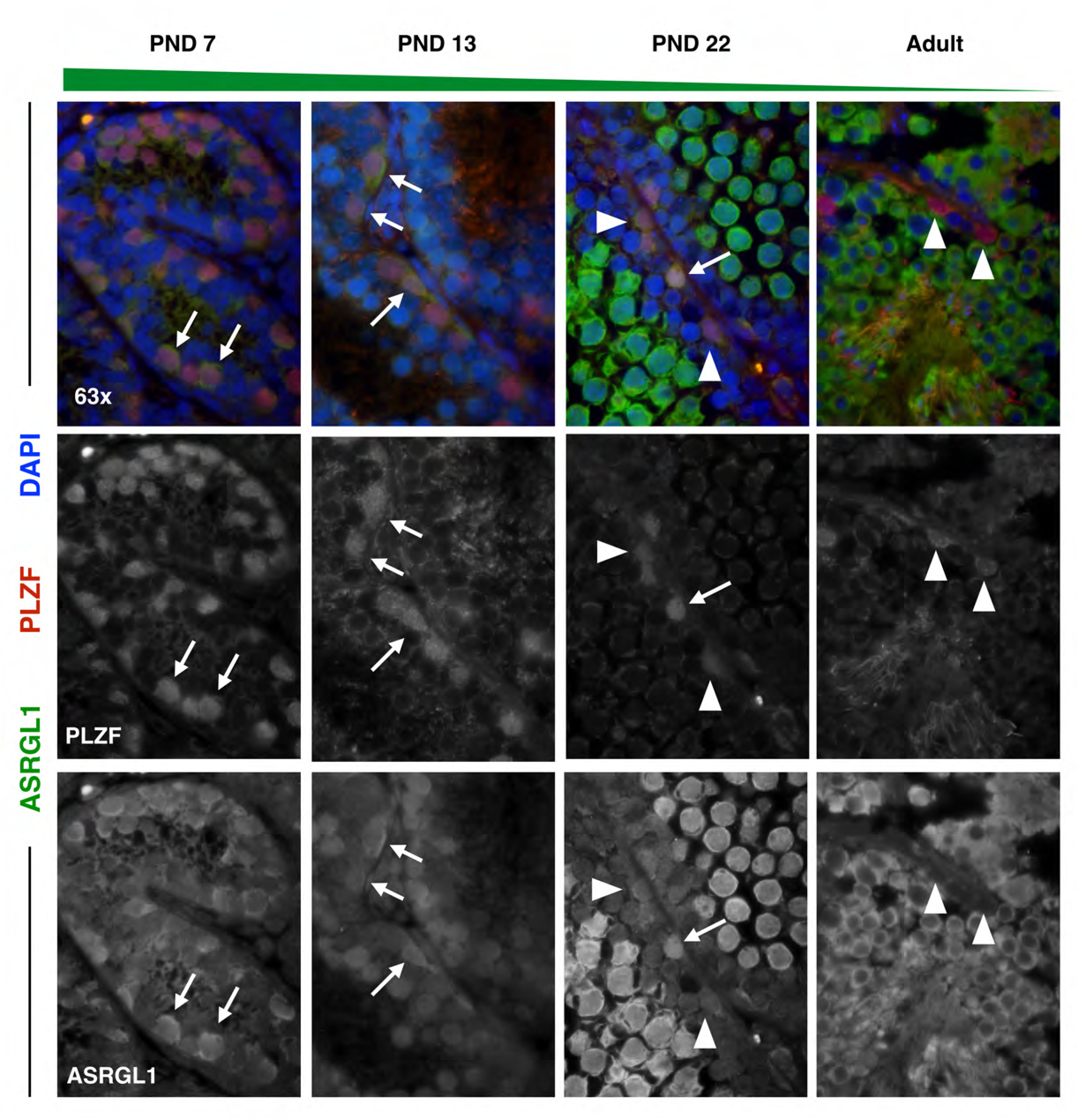
Split channels of ARSGL1 expression in spermatogonia. Spermatogonial marker PLZF (red) and ASRGL1 (green) were stained in 5μm testis tissue sections from mice ages PND7, PND13, PND22, and adult. DAPI (blue) denotes nuclei. ASRGL1 protein expression decreases in PLZF+ spermatogonia with age. High-ASRGL1-expressing spermatogonia are indicated by full arrows with a line, while low-ASRGL1-expressing spermatogonia are indicated by arrowheads. Individual channels are represented in gray scale.

**Supplementary Figure 8.**
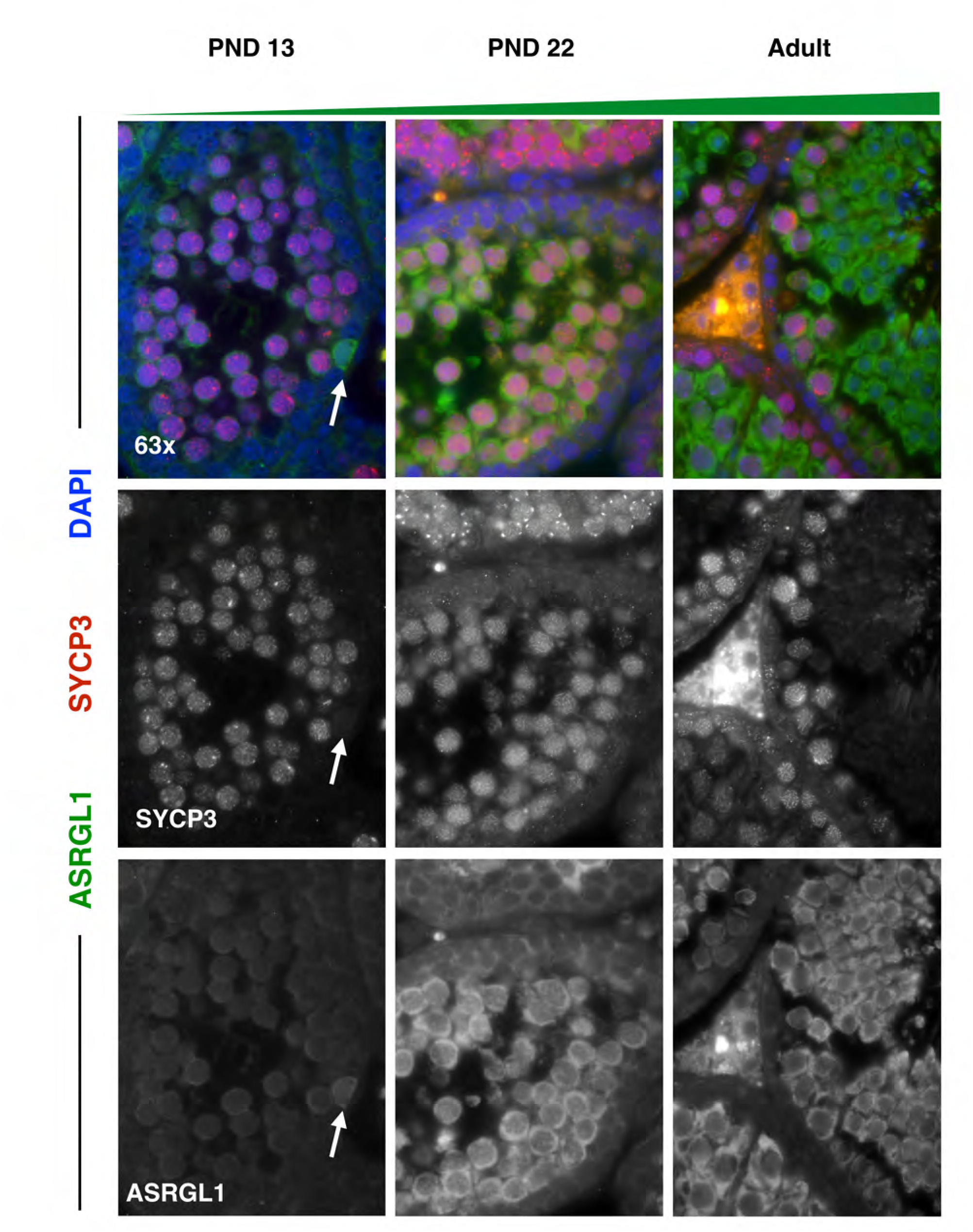
Split channels of ARSGL1 expression in spermatocytes. Spermatocyte marker SYCP3 (red) and ASRGL1 (green) were stained in 5μm testis tissue sections from mice ages PND13, PND22, and adult. ASRGL1 protein expression increases in SYCP3+ spermatocytes with age. For all images, high-ASRGL1-expressing spermatogonia are indicated by full arrows with a line. Individual channels are represented in gray scale.

**Supplementary Figure 9.**
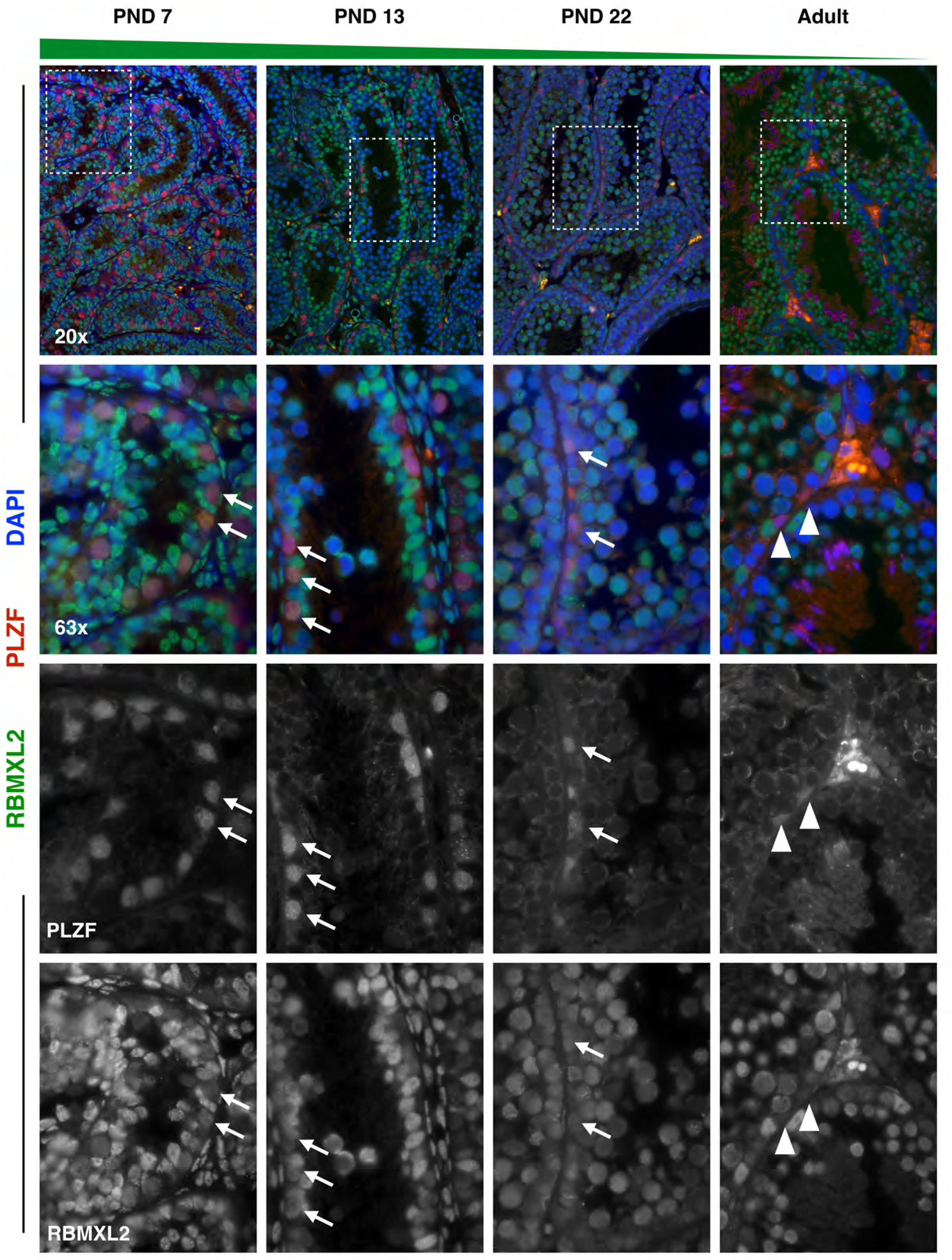
RBMXL2 is highly expressed specifically in spermatogonia from neonatal and juvenile mice, decreasing with age. Spermatogonial marker PLZF (red) and RBMXl2 (green) were stained in 5μm testis tissue sections from mice ages PND7, PND13, PND22, and adult. DAPI (blue) denotes nuclei. RBMXL2 protein expression decreases in PLZF+ spermatogonia with age. High-RBMXL2-expressing spermatogonia are indicated by full arrows with a line, while low-RBMXL2-expressing cells are indicated by arrowheads. Individual channels are represented in gray scale.

**Supplementary Figure 10.**
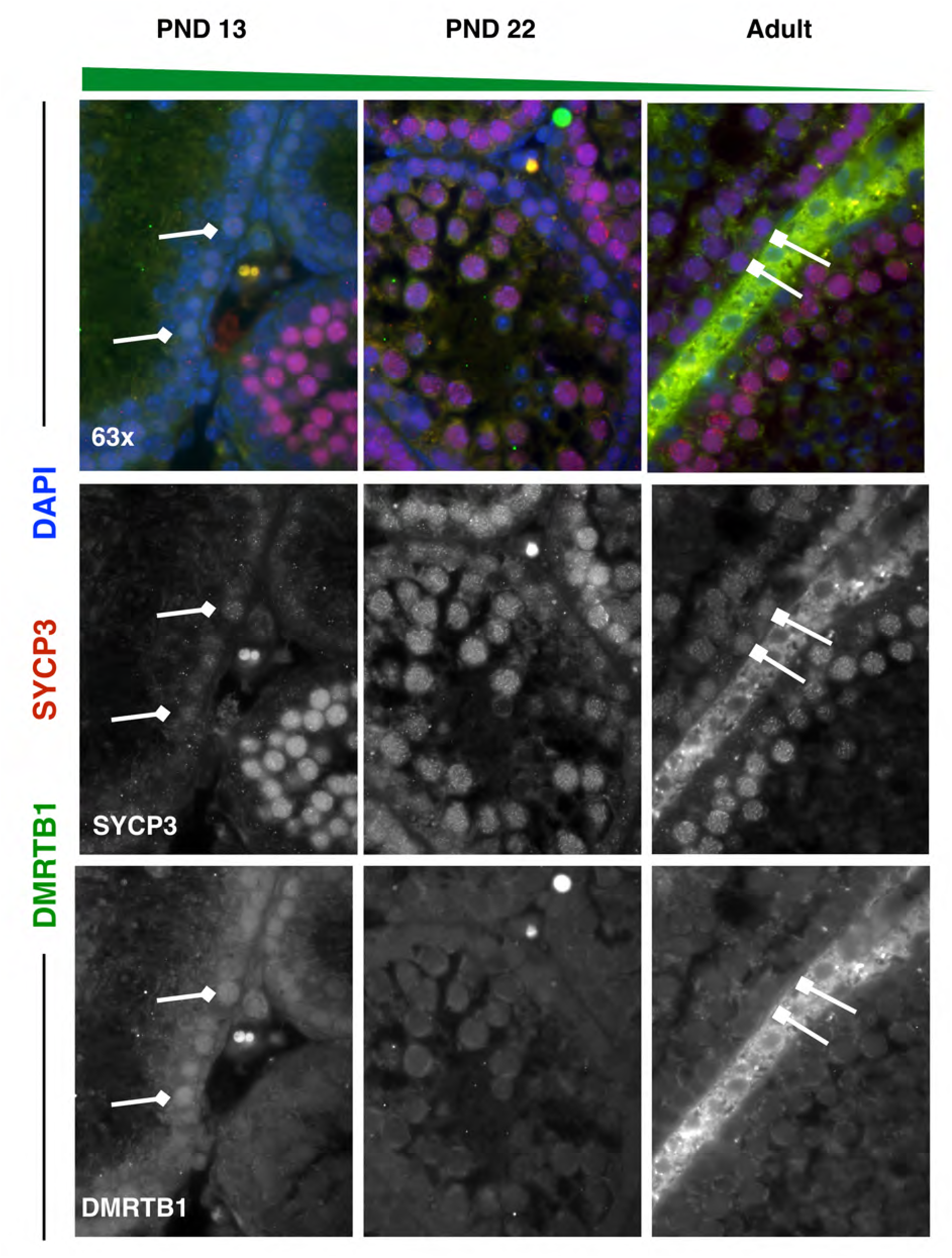
Split channels of DMRTB1 expression in spermatocytes. Spermatocyte marker SYCP3 (red) and DMRTB1 (green) were stained in 5μm testis tissue sections from mice ages PND13, PND22, and adult. DMRTB1 protein is expressed in the nucleus of first-wave spermatocytes at PND13, with decreasing expression in pachyene spermatocytes with age. For all images, high-DMRTB1-expressing spermatocytes are indicated by diamond-headed arrows, while low-DMRTB1-expressing spermatocytes are indicated by square-headed arrows. Individual channels are represented in gray scale.

**Supplementary Figure 11.**
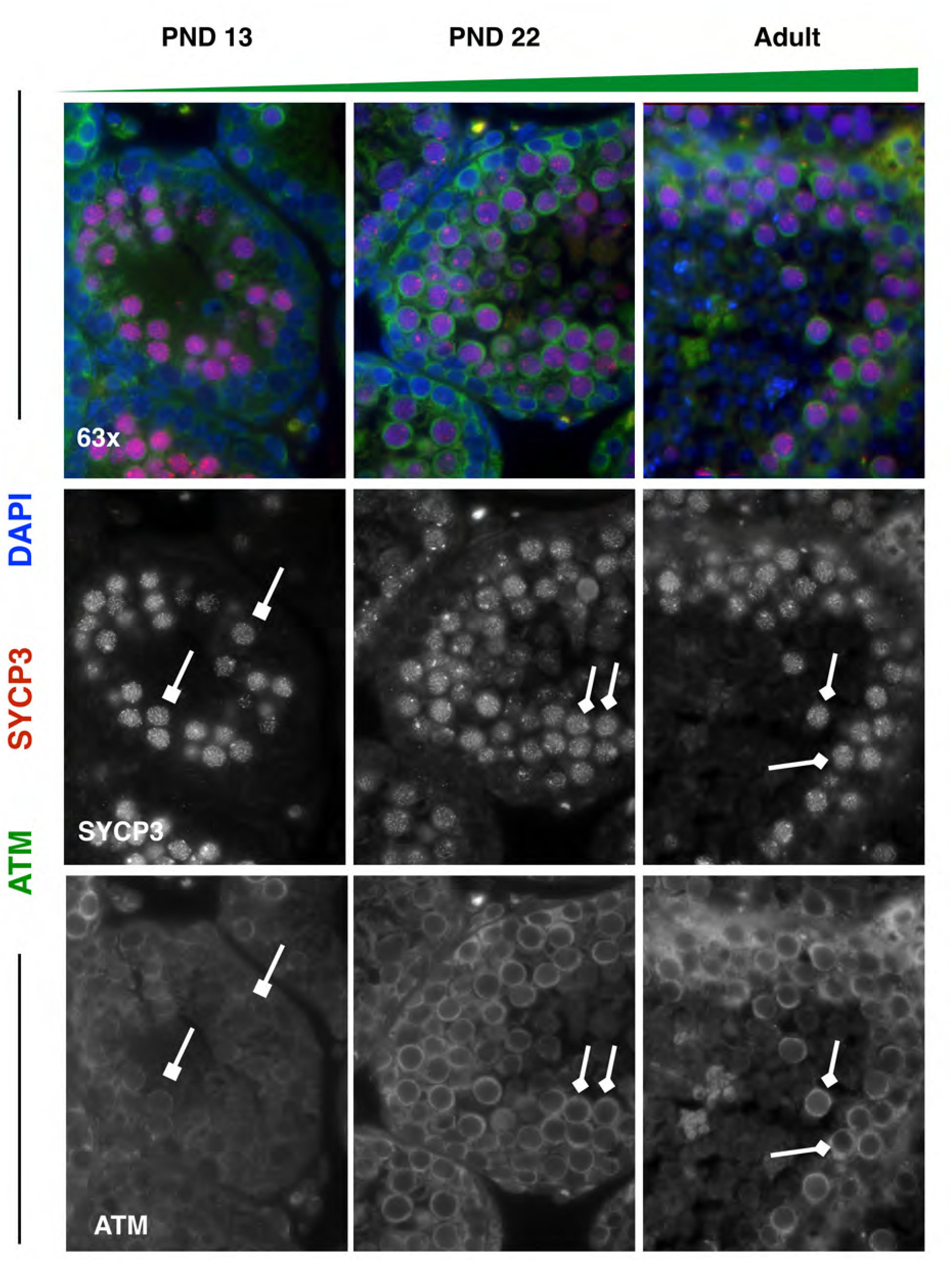
Split channels of ATM expression in spermatocytes. Spermatocyte marker SYCP3 (red) and ATM (green) were stained in 5μm testis tissue sections from mice ages PND13, PND22, and adult. ATM protein expression increases in SYCP3+ spermatocytes with age. For all images, high-ATM-expressing spermatocytes are indicated by diamond-headed arrows, while low-ATM-expressing spermatocytes are indicated by square-headed arrows. Individual channels are represented in gray scale.

**Supplementary Figure 12.**
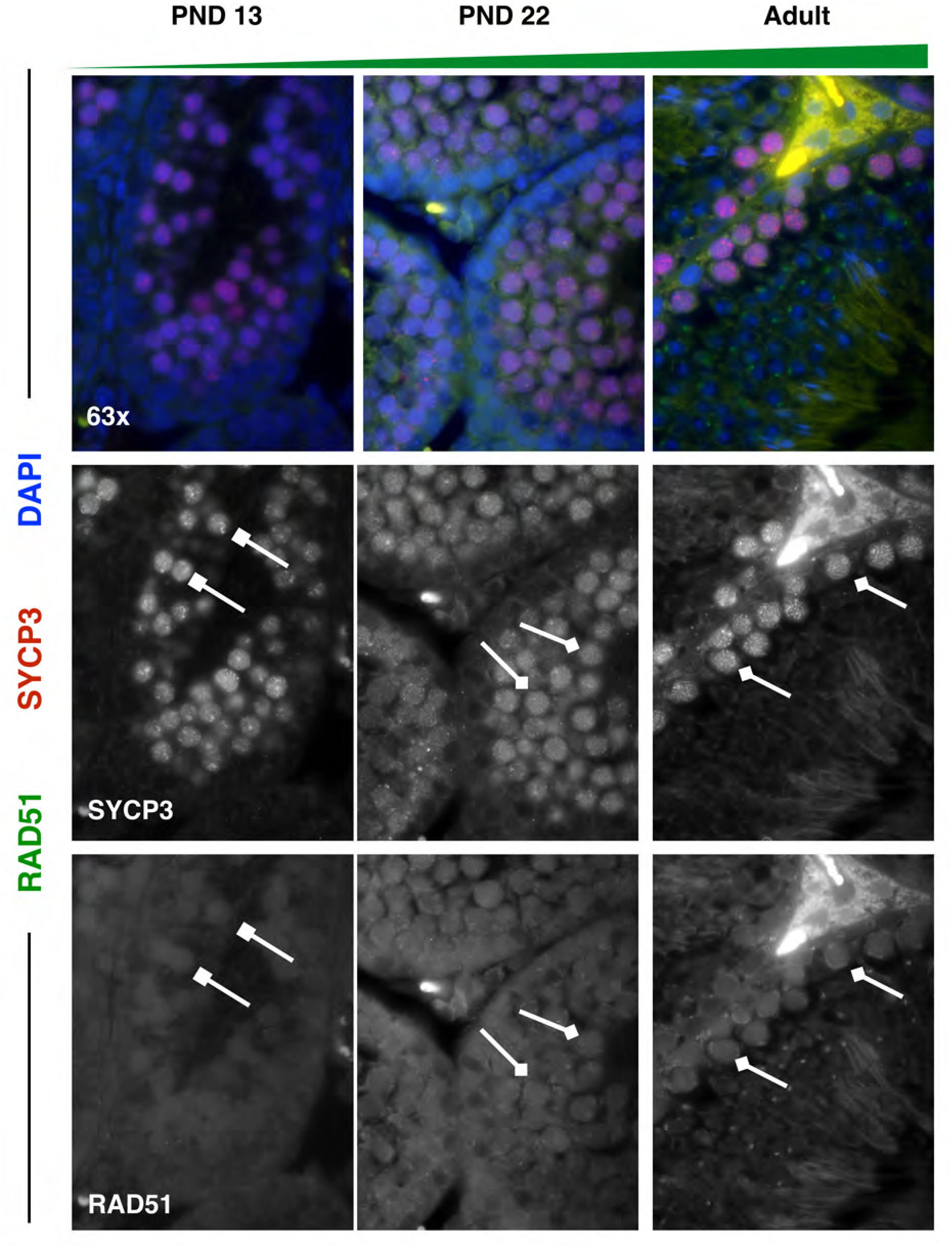
Split channels of RAD51 expression in spermatocytes. Spermatocyte marker SYCP3 (red) and RAD51 (green) were stained in 5μm testis tissue sections from mice ages PND13, PND22, and adult. RAD51 protein expression increases in SYCP3+ spermatocytes with age. For all images, high-RAD51-expressing spermatocytes are indicated by diamond-headed arrows, while low-RAD51-expressing spermatocytes are indicated by square-headed arrows. Individual channels are represented in gray scale.

**Table S1** - *Genes expressed in specific germ cell types*. Marker genes were determined that distinguish different primary cell clusters. Up to 250 genes per primary cell type are listed, with statistics from Seurat comparing expression in the marker-associated cell type (X.1) to all other germ cells (X.2). Data for these genes are depicted as row-normalized gene expression in individual cells in **Figure 3**.

**Table S2** - *Genes with high counts in spermatids (filtered out)*. Due to contaminating cell-free mRNA derived from lysed spermatids (detected only in samples in which spermatids are present), these genes expressed at high levels in spermatids were removed from the dataset. The UMI1/UMI2 ratio reflects the expression of each gene in spermatids relative to all other germ cells.

**Table S3** - *Genes with variable expression in spermatogonia during testis maturation.* Model-based analysis of single-cell transcriptomics (MAST) was utilized to identify genes that are variably expressed in spermatogonia as a function of mouse age. Genes listed in this table include “markers” (genes with significantly upregulated expression in all spermatogonia), as well as genes with significant variation in expression in spermatogonia during testis maturation. Data for these genes are depicted as row-normalized gene expression in individual cells in **Figure 4**.

**Table S4** - *Gene Set Enrichment Analysis of Reactome pathways in spermatogonia during testis maturation*. Reactome pathways with significantly differential expression as a function of mouse age in spermatogonia are included (FDR < 0.05); the same results are displayed in Cytoscape diagrams in **Figure 5**. The table includes GSEA results for each significant Reactome pathway, including the gene list, enrichment score (ES), normalized enrichment score (NES), and results of the statistical test for enrichment.

**Table S5** - *Genes with variable expression in spermatocytes during testis maturation.* Model-based analysis of single-cell transcriptomics (MAST) was utilized to identify genes that are variably expressed in spermatocytes as a function of age. Genes listed in this table include “markers” (genes with significantly upregulated expression in all spermatocytes), as well as genes with significant variation in expression in spermatocytes during testis maturation. Data for these genes are depicted as row-normalized gene expression in individual cells in **Figure 6**.

**Table S6** - *Gene Set Enrichment Analysis of Reactome pathways in spermatocytes during testis maturation.* Reactome pathways with significant differential expression as a function of mouse age in spermatocytes are included (FDR < 0.05); the same results are displayed in Cytoscape diagrams in **Figure 7**. The table includes GSEA results for each significant Reactome pathway, including the gene list, enrichment score (ES), normalized enrichment score (NES), and results of the statistical test for enrichment.

**Table S7** – *Antibodies used for immunofluorescent staining*. All primary and secondary antibodies used for immunofluorescence validation are documented with product numbers and dilutions used.

## References

1. Phillips, B. T., Gassei, K. & Orwig, K. E. Spermatogonial stem cell regulation and spermatogenesis. Philos. Trans. R. Soc. Lond. B. Biol. Sci. 365, 1663–78 (2010).

2. Lawson, K. A. & Hage, W. J. *Germline development*. *Ciba Foundation Symposium* 182, (Wiley, 1994).

3. Anderson, R., Copeland, T. K., Schöler, H., Heasman, J. & Wylie, C. The onset of germ cell migration in the mouse embryo. Mech. Dev. 91, 61–68 (2000).

4. Nakatsuji, N. & Chuma, S. Differentiation of mouse primordial germ cells into female or male germ cells. Int. J. Dev. Biol. 45, 541–8 (2001).

5. Ewen, K. A. & Koopman, P. Mouse germ cell development: From specification to sex determination. Molecular and Cellular Endocrinology 323, 76–93 (2010).

6. Yoshida, S. et al. The first round of mouse spermatogenesis is a distinctive program that lacks the self-renewing spermatogonia stage. Development 133, 1495–1505 (2006).

7. Tegelenbosch, R. & de Rooij, D. A quantitative study of spermatogonial multiplication and stem cell renewal in the C3H/101 F1 hybrid mouse. Mutat. Res. 290, 193–200 (1993).

8. Koubova, J. et al. Retinoic acid regulates sex-specific timing of meiotic initiation in mice. Proc. Natl. Acad. Sci. 103, 2474–2479 (2006).

9. Anderson, E. L. et al. Stra8 and its inducer, retinoic acid, regulate meiotic initiation in both spermatogenesis and oogenesis in mice. Proc. Natl. Acad. Sci. U. S. A. 105, 14976–14980 (2008).

10. Endo, T. et al. Periodic retinoic acid–STRA8 signaling intersects with periodic germ-cell competencies to regulate spermatogenesis. Proc. Natl. Acad. Sci. 112, E2347–E2356 (2015).

11. Endo, T., Freinkman, E., de Rooij, D. G. & Page, D. C. Periodic production of retinoic acid by meiotic and somatic cells coordinates four transitions in mouse spermatogenesis. Proc. Natl. Acad. Sci. 114, E10132–E10141 (2017).

12. Baarends, W. M. et al. Chromatin dynamics in the male meiotic prophase. Cytogenet. Genome Res. 103, 225–34 (2003).

13. Gray, S. & Cohen, P. E. Control of Meiotic Crossovers: From Double-Strand Break Formation to Designation. Annu. Rev. Genet. 50, 175–210 (2016).

14. Vrooman, L. A., Nagaoka, S. I., Hassold, T. J. & Hunt, P. A. Evidence for paternal age-related alterations in meiotic chromosome dynamics in the mouse. Genetics 196, 385–96 (2014).

15. Zelazowski, M. J., Sandoval, M. & Gribbell, M. A. Age-Dependent Alterations in Meiotic Recombination Cause Chromosome Segregation Errors in Spermatocytes In Brief. Cell 171, 601–607.e13 (2017).

16. Goodyear, S. & Brinster, R. Isolation of the Spermatogonial Stem Cell-Containing Fraction from Testes. Cold Spring Harb. Protoc. 2017, pdb.prot094185 (2017).

17. Osuru, H. P. et al. The acrosomal protein SP-10 (*Acrv1*) is an ideal marker for staging of the cycle of seminiferous epithelium in the mouse. Mol. Reprod. Dev. 81, 896–907 (2014).

18. Zheng, G. X. Y. et al. Massively parallel digital transcriptional profiling of single cells. Nat. Commun. 8, 14049 (2017).

19. Finak, G. et al. MAST: a flexible statistical framework for assessing transcriptional changes and characterizing heterogeneity in single-cell RNA sequencing data. Genome Biol. 16, 278 (2015).

20. Subramanian, A. et al. Gene set enrichment analysis: a knowledge-based approach for interpreting genome-wide expression profiles. Proc. Natl. Acad. Sci. U. S. A. 102, 15545–50 (2005).

21. Shannon, P. et al. Cytoscape: a software environment for integrated models of biomolecular interaction networks. Genome Res. 13, 2498–504 (2003).

22. Merico, D., Isserlin, R., Stueker, O., Emili, A. & Bader, G. D. Enrichment map: a network-based method for gene-set enrichment visualization and interpretation. PLoS One 5, e13984 (2010).

23. Kubota, H., Avarbock, M. R. & Brinster, R. L. Spermatogonial stem cells share some, but not all, phenotypic and functional characteristics with other stem cells. Proc. Natl. Acad. Sci. 100, 6487–6492 (2003).

24. Schrans-Stassen, B. H. G. J., van de Kant, H. J. G., de Rooij, D. G. & van Pelt, A. M. M. Differential Expression of c-*kit* in Mouse Undifferentiated and Differentiating Type A Spermatogonia. Endocrinology 140, 5894–5900 (1999).

25. Busada, J. T., Niedenberger, B. A., Velte, E. K., Keiper, B. D. & Geyer, C. B. Mammalian target of rapamycin complex 1 (mTORC1) Is required for mouse spermatogonial differentiation in vivo. Dev. Biol. 407, 90–102 (2015).

26. Vincent, S. et al. Stage-specific expression of the Kit receptor and its ligand (KL) during male gametogenesis in the mouse: a Kit-KL interaction critical for meiosis. Development 125, 4585–93 (1998).

27. Pui, H. P. & Saga, Y. Gonocytes-to-spermatogonia transition initiates prior to birth in murine testes and it requires FGF signaling. Mech. Dev. 144, 125–139 (2017).

28. Hasegawa, K. & Saga, Y. FGF8-FGFR1 Signaling Acts as a Niche Factor for Maintaining Undifferentiated Spermatogonia in the Mouse1. Biol. Reprod. 91, 145 (2014).

29. Sun, X. et al. FancJ (Brip1) loss-of-function allele results in spermatogonial cell depletion during embryogenesis and altered processing of crossover sites during meiotic prophase I in mice. Chromosoma 125, 237–52 (2016).

30. Blackshear, P. E. et al. Brca1 and Brca2 expression patterns in mitotic and meiotic cells of mice. Oncogene 16, 61–68 (1998).

31. Simhadri, S. et al. Male fertility defect associated with disrupted BRCA1-PALB2 interaction in mice. J. Biol. Chem. 289, 24617–29 (2014).

32. Broering, T. J. et al. BRCA1 establishes DNA damage signaling and pericentric heterochromatin of the X chromosome in male meiosis. J. Cell Biol. 205, 663–75 (2014).

33. Haaf, T., Golub, E. I., Reddy, G., Radding, C. M. & Ward, D. C. Nuclear foci of mammalian Rad51 recombination protein in somatic cells after DNA damage and its localization in synaptonemal complexes. Proc. Natl. Acad. Sci. U. S. A. 92, 2298–302 (1995).

34. Testa, E. et al. H2AFX and MDC1 promote maintenance of genomic integrity in male germ cells. J. Cell Sci. 131, jcs214411 (2018).

35. Plug, A. W. et al. ATM and RPA in meiotic chromosome synapsis and recombination. Nat. Genet. 17, 457–461 (1997).

36. Costoya, J. A. et al. Essential role of Plzf in maintenance of spermatogonial stem cells. Nat. Genet. 36, 653–659 (2004).

37. Filipponi, D. et al. Repression of kit Expression by Plzf in Germ Cells. Mol. Cell. Biol. 27, 6770–6781 (2007).

38. Lovelace, D. L. et al. The regulatory repertoire of PLZF and SALL4 in undifferentiated spermatogonia. Development 143, 1893–1906 (2016).

39. Yuan, L. et al. The murine SCP3 gene is required for synaptonemal complex assembly, chromosome synapsis, and male fertility. Mol. Cell 5, 73–83 (2000).

40. Cantor, J. R., Stone, E. M., Chantranupong, L. & Georgiou, G. The Human Asparaginase-like Protein 1 hASRGL1 Is an Ntn Hydrolase with β-Aspartyl Peptidase Activity. Biochemistry 48, 11026–11031 (2009).

41. Biswas, P. et al. A missense mutation in *ASRGL1* is involved in causing autosomal recessive retinal degeneration. Hum. Mol. Genet. 25, ddw113 (2016).

42. Edqvist, P.-H. D. et al. Loss of ASRGL1 expression is an independent biomarker for disease-specific survival in endometrioid endometrial carcinoma. Gynecol. Oncol. 137, 529–537 (2015).

43. Fonnes, T. et al. Asparaginase-like protein 1 expression in curettage independently predicts lymph node metastasis in endometrial carcinoma: a multicenter study. BJOG An Int. J. Obstet. Gynaecol. (2018). doi:10.1111/1471-0528.15403

44. Huvila, J. et al. Combined ASRGL1 and p53 immunohistochemistry as an independent predictor of survival in endometrioid endometrial carcinoma. Gynecol. Oncol. 149, 173–180 (2018).

45. Fonnes, T. et al. Asparaginase-like protein 1 is an independent prognostic marker in primary endometrial cancer, and is frequently lost in metastatic lesions. Gynecol. Oncol. 148, 197–203 (2018).

46. Bush, L. A. et al. A novel asparaginase-like protein is a sperm autoantigen in rats. Mol. Reprod. Dev. 62, 233–247 (2002).

47. Elliott, D. J. et al. An evolutionarily conserved germ cell-specific hnRNP is encoded by a retrotransposed gene. Hum. Mol. Genet. 9, 2117–2124 (2000).

48. Ehrmann, I. et al. Haploinsufficiency of the germ cell-specific nuclear RNA binding protein hnRNP G-T prevents functional spermatogenesis in the mouse. Hum. Mol. Genet. 17, 2803–2818 (2008).

49. Maymon, B. B.-S. et al. Localization of the germ cell-specific protein, hnRNP G-T, in testicular biopsies of azoospermic men. Acta Histochem. 104, 255–261 (2002).

50. Zhang, T., Murphy, M. W., Gearhart, M. D., Bardwell, V. J. & Zarkower, D. The mammalian Doublesex homolog DMRT6 coordinates the transition between mitotic and meiotic developmental programs during spermatogenesis. Development 141, 3662–3671 (2014).

51. Hamer, G., Kal, H. B., Westphal, C. H., Ashley, T. & de Rooij, D. G. Ataxia Telangiectasia Mutated Expression and Activation in the Testis1. Biol. Reprod. 70, 1206–1212 (2004).

52. Meng, X. et al. Regulation of cell fate decision of undifferentiated spermatogonia by GDNF. Science 287, 1489–93 (2000).

53. Hofmann, M.-C., Braydich-Stolle, L. & Dym, M. Isolation of male germ-line stem cells; influence of GDNF. Dev. Biol. 279, 114–124 (2005).

54. Kubota, H., Avarbock, M. R. & Brinster, R. L. Growth factors essential for self-renewal and expansion of mouse spermatogonial stem cells. Proc. Natl. Acad. Sci. 101, 16489–16494 (2004).

55. Busada, J. T. et al. Retinoic acid regulates Kit translation during spermatogonial differentiation in the mouse. Dev. Biol. 397, 140–149 (2015).

56. Gonzalez-Herrera, I. G. et al. Testosterone regulates FGF-2 expression during testis maturation by an IRES-dependent translational mechanism. FASEB J. 20, 476–8 (2006).

57. Roecker, G. O. & Huether, C. A. An analysis for paternal-age effect in Ohio’s Down syndrome births, 1970-1980. Am. J. Hum. Genet. 35, 1297–306 (1983).

58. Steiner, B. et al. An unexpected finding: younger fathers have a higher risk for offspring with chromosomal aneuploidies. Eur. J. Hum. Genet. 23, 466–72 (2015).

59. Lange, J. et al. ATM controls meiotic double-strand-break formation. Nature 479, 237–240 (2011).

60. Yoshida, S., Nabeshima, Y.-I. & Nakagawa, T. Stem Cell Heterogeneity: Actual and Potential Stem Cell Compartments in Mouse Spermatogenesis. Ann. N. Y. Acad. Sci. 1120, 47–58 (2007).

61. Zheng, K., Wu, X., Kaestner, K. H. & Wang, P. The pluripotency factor LIN28 marks undifferentiated spermatogonia in mouse. BMC Dev. Biol. 9, 38 (2009).

62. Hermann, B. P., Phillips, B. T. & Orwig, K. E. The Elusive Spermatogonial Stem Cell Marker?1. Biol. Reprod. 85, 221–223 (2011).

63. Gassei, K. & Orwig, K. E. SALL4 Expression in Gonocytes and Spermatogonial Clones of Postnatal Mouse Testes. PLoS One 8, e53976 (2013).

64. Sablitzky, F. et al. Stage-and subcellular-specific expression of Id proteins in male germ and Sertoli cells implicates distinctive regulatory roles for Id proteins during meiosis, spermatogenesis, and Sertoli cell function. Cell Growth Differ. 9, 1015–24 (1998).

65. Oatley, M. J., Kaucher, A. V., Racicot, K. E. & Oatley, J. M. Inhibitor of DNA Binding 4 Is Expressed Selectively by Single Spermatogonia in the Male Germline and Regulates the Self-Renewal of Spermatogonial Stem Cells in Mice1. Biol. Reprod. 85, 347–356 (2011).

66. Sun, F., Xu, Q., Zhao, D. & Degui Chen, C. Id4 Marks Spermatogonial Stem Cells in the Mouse Testis. Sci. Rep. 5, 17594 (2015).

67. Morimoto, H. et al. Phenotypic Plasticity of Mouse Spermatogonial Stem Cells. PLoS One 4, e7909 (2009).

68. Cooke, P. S., Simon, L., Nanjappa, M. K., Medrano, T. I. & Berry, S. E. Plasticity of spermatogonial stem cells. Asian J. Androl. 17, 355–9

69. Butler, A., Hoffman, P., Smibert, P., Papalexi, E. & Satija, R. Integrating single-cell transcriptomic data across different conditions, technologies, and species. Nat. Biotechnol. 36, 411–420 (2018).

70. Johnson, W. E., Li, C. & Rabinovic, A. Adjusting batch effects in microarray expression data using empirical Bayes methods. Biostatistics 8, 118–27 (2007).

71. Soumillon, M. et al. Cellular source and mechanisms of high transcriptome complexity in the mammalian testis. Cell Rep. 3, 2179–90 (2013).

72. Min Jung, Daniel Wells, Janette Rusch, Suhaira Ahmed, Jonathan Marchini, Simon Myers, D. C. Unified single-cell analysis of testis gene regulation and pathology in 5 mouse strains. BioRxiv (2018).

73. von Kopylow, K. & Spiess, A.-N. Human spermatogonial markers. Stem Cell Res. 25, 300–309 (2017).

